# No strong evidence that social network index is associated with gray matter volume from a data-driven investigation

**DOI:** 10.1101/2019.12.19.883173

**Authors:** Chujun Lin, Umit Keles, J. Michael Tyszka, Marcos Gallo, Lynn Paul, Ralph Adolphs

**Affiliations:** Division of Humanities and Social Sciences, California Institute of Technology, CA, USA; Division of Biology and Biological Engineering, California Institute of Technology, CA, USA

**Keywords:** social network index, gray matter volume, predictive modeling, cross-validation

## Abstract

Recent studies in adult humans have reported correlations between individual differences in people’s Social Network Index (SNI) and gray matter volume (GMV) across multiple regions of the brain. However, the cortical and subcortical loci identified are inconsistent across studies. These discrepancies might arise because different regions of interest were hypothesized and tested in different studies without controlling for multiple comparisons, and/or from insufficiently large sample sizes to fully protect against statistically unreliable findings. Here we took a data-driven approach in a pre-registered study to comprehensively investigate the relationship between SNI and GMV in every cortical and subcortical region, using three predictive modeling frameworks. We also included psychological predictors such as cognitive and emotional intelligence, personality, and mood. In a sample of healthy adults (n = 92), neither multivariate frameworks (e.g., ridge regression with cross-validation) nor univariate frameworks (e.g., univariate linear regression with cross-validation) showed a significant association between SNI and any GMV or psychological feature after multiple comparison corrections (all R-squared values ≤ 0.1). These results emphasize the importance of large sample sizes and hypothesis-driven studies to derive statistically reliable conclusions, and suggest that future meta-analyses will be needed to more accurately estimate the true effect sizes in this field.

## 1. Introduction

It has been well-documented that neocortex volume is positively correlated with social group size across multiple primate species (Dunbar, 1998; Dunbar & Shultz, 2007), an intriguing finding that has motivated a number of subsequent studies in humans (see below). It is important to keep in mind that social group size is of course not the only factor in the evolution of large brains: it is merely one variable amongst many interacting variables that determines fitness. For instance, diet and other ecological variables are also associated with brain size (Barton, 1999). Nonetheless, across the many variables that contribute to brain size (or to gray matter volume of specific structures), social group size remains as one of the most robust when studies examine this question across species (Dunbar & Shultz, 2017).

While the correlation between brain volume and social group size is robust across species, it has also been suggested that a similar association might obtain across individuals within a species: some individuals are embedded in larger or smaller social groups, and one might expect this variation in social behavior to be related to the brain. In particular, one might expect the variation to be related to brain structures implicated in social cognition. A number of studies have examined this within-species hypothesis in humans (Table 1) by correlating GMV of structures such as amygdala with various social network metrics, in particular self-reports of the number of people one has contacted within a given period, such as the social network index or SNI, a metric we also used in the present study.

**Table 1.** Summary of previous studies in humans on the correlations between social network metrics and GMV of cortical and subcortical structures of the brain. Abbreviations: L left, R right, ITS inferior temporal sulcus, SFG superior frontal gyrus, ACC anterior cingulate cortex, mPFC medial prefrontal cortex, TPJ temporoparietal junction, STS superior temporal sulcus, OFC orbitofrontal cortex, AIC anterior insular cortex.

| Literature | Hypothesized Regions (ROIs) | Social Network Metrics | Sample Size | Age Range | Correction for Multiple Comparisons | Significant Regions |
| --- | --- | --- | --- | --- | --- | --- |
| Bickart et al., 2011 | 1) amygdala<br>2) hippocampus<br>3) exploratory analysis of all other subcortical regions<br>4) exploratory analysis of all cortical thickness | 2 subscales of SNI: the number of people in social network, the number of embedded networks | N = 58 | 19 - 83 | For 1) and 2): linear regressions, uncorrected<br>For 3): linear regressions, Bonferroni correction for testing multiple regions, but not for multiple SNIs<br>For 4): general linear regressions, uncorrected | L amygdala<br>R amygdala<br>If uncorrected for multiple comparison ($p < 0.01$), also:<br>R subgenual ACC<br>L caudal SFG<br>L caudal ITS |
| Lewis et al., 2011 | 1) mPFC<br>2) TPJ<br>3) STS<br>4) frontal pole | Dunbar's number: the number of people contacted in previous 30 days | N = 45 | 18 - 50 | $p < 0.001$ uncorrected with an extent threshold of $>5$ voxels within ROIs<br>*survived small volume correction at $p = 0.05$ with 8mm radius spheres | *Ventromedial frontal gyrus<br>Medial orbitofrontal gyrus |
| Kanai et al., 2012 | 1) amygdala<br>2) posterior STS<br>3) TPJ<br>4) mPFC<br>5) precuneus<br>6) medial temporal lobe | Online social network size: the number of Facebook friends | Sample 1<br>N = 125<br>Sample 2<br>N = 40 | Sample 1<br>$23 \pm 4$<br>Sample 2<br>$22 \pm 3$ | Sample 1<br>$p < 0.05$ family-wise error corrected for the whole-brain volume<br>*only survived correction for small volumes of 10 mm spheres around ROIs<br>Sample 2<br>$p < 0.05$ uncorrected for testing multiple loci identified in Sample 1 | R posterior STS<br>R entorhinal<br>L middle temporal gyrus<br>*L amygdala<br>*R amygdala |
| Bickart et al., 2012 | 1) amygdala, controlling for network connectivity | All three subscales of SNI | N = 29 | 19 - 32 | Linear regressions, uncorrected | Amygdala |
| Powell, et al., 2012 | 1) orbital PFC<br>2) dorsal PFC | The number of people contacted in previous 7 days | N = 40 | 18 - 47 | Path analysis, uncorrected | Orbital PFC |
| Von Der Heide et al., 2014 | 1) amygdala<br>2) R subgenual ACC<br>3) L posterior ITS<br>4) L posterior SFG<br>5) posterior STS<br>6) middle temporal gyrus<br>7) entorhinal<br>8) OFC | 3 measures: the number of Facebook friends, Dunbar's number, Norbeck Social Support | N = 40 females | 12 - 30 | $p < 0.05$ family-wise error corrected for small volumes of 10 mm radius spheres around the ROIs; uncorrected for testing multiple measures | L amygdala,<br>R amygdala,<br>L posterior ITS<br>L posterior SFG<br>L entorhinal<br>R entorhinal<br>L OFC<br>R OFC |
| Noonan et al., 2018 | 1) ACC | 2 measures: the number of people contacted in | N = 18 | $52 \pm 15$ | Whole brain approach only reporting regions that bilaterally survive ($p < 0.0001$) with an | Subcallosal parts of vmPFC<br>Anterior temporal cortex |
| | | previous 7 days, the number of people contacted in previous 30 days | | | extent threshold of >40 voxels<br><u>ROI approach</u><br>$p < 0.05$ family-wise error corrected for small volumes of all voxels in the ROI | The border of posterior cingulate cortex and precuneus |
| Spagna et al., 2018 | 1) AIC<br>2) amygdala<br>3) exploratory analysis of all other cortical thickness | A composite measure of all three subscales of SNI | Sample 1<br>N = 50<br>Sample 2<br>N = 100 | Sample 1<br>19 - 37<br>Sample 2<br>18 - 29 | For 1) and 2): $p < 0.05$ with contiguous-voxel extent thresholds estimated using AlphaSim<br>For 3): $p < 0.05$ family-wise error corrected for the whole-brain volume | L AIC in Sample 1<br>R AIC in Sample 2<br>and when both samples were combined |
| Kwak et al., 2018 | 1) amygdala<br>2) OFC<br>3) dorsal mPFC<br>4) TPJ<br>5) precuneus | The number of people discussed things with in the last 12 months | N = 68 | 59 - 84 | $p < 0.05$ family-wise error corrected with a cluster defining threshold of $p < 0.001$ estimated by the Gaussian random field | R OFC<br>dorsal mPFC |

A study in macaques even suggests the causal hypothesis that social group size could cause changes in brain size (Sallet et al., 2011): macaques randomly assigned to live in larger groups showed increased GMV in certain brain structures thought to underlie social cognition. Whether on the timescale of evolution or of the life of an individual, the above varied findings raise the hypothesis that social network metrics in humans might be correlated with GMV in specific brain structures.

However, characterizing social networks in humans is fundamentally different from quantifying social group size in other primates due to the greater complexity and variability of human social relationships (Dunbar, 1998). Previous studies attempting to test the within-species hypothesis in humans (Table 1) have employed various metrics of social networks, such as the number of people one had seen or talked to at least once every two weeks (Bickart, Hollenbeck, Barrett, & Dickerson, 2012; Bickart, Wright, Dautoff, Dickerson, & Barrett, 2011; Bickart et al., 2011), the number of people one had contacted over the last 12 months, 30 days, or 7 days (Kwak, Joo, Youm, & Chey, 2018; Lewis, Rezaie, Brown, Roberts, & Dunbar, 2011; Noonan, Mars, Sallet, Dunbar, & Fellows, 2018; Powell Joanne, Lewis Penelope A., Roberts Neil, García-Fiñana Marta, & Dunbar R. I. M., 2012), or the number of friends one had on social media (Kanai, Bahrami, Roylance, & Rees, 2012). While all those metrics can fluctuate over months, weeks, and even days for an individual, GMV of brain structures are relatively stable over time in healthy adults. This makes at least some metrics of social networks in humans, such as the SNI, prima facie implausible candidates for being correlated with variability in structural brain measures, raising some caution about how to interpret any putative findings.

Indeed, previous studies in humans investigating the relationship between social network metrics and GMV have produced inconsistent results (Table 1). For instance, while some studies showed that bilateral amygdala volume was positively correlated with SNI (Bickart et al., 2011), others failed to replicate these relationships (Spagna et al., 2018). In addition, the different regions of interest hypothesized, and different methods for correcting for multiple comparisons used in past research might also contribute to the discrepant findings (Kanai et al., 2012; Lewis et al., 2011; Noonan et al., 2018).

Here, we took a purely data-driven approach to examine the relationship between SNI and GMV, with the aim of uncovering any relationships with specific brain regions. We did not hypothesize SNI to correlate with GMV of any specific brain region, and instead comprehensively tested the effect of every cortical and subcortical volume to see if an agnostic approach would discover (or reproduce) any candidates. We examined these relationships using three different predictive modeling frameworks, which capitalized on the strengths of both multivariate analyses and univariate analyses, explored the prediction performance with or without feature selection, and implemented cross-validation to increase the generalizability of our results. To handle multiple comparisons, all effects within a framework was corrected for false discovery rate (FDR). Since previous studies have also reported that various psychological measures such as personality and perceived stress were linked to individual differences in social networks (Asendorpf & Wilpers, 1998; Nabi, Prestin, & So, 2013), we also included a list of psychological measures in our frameworks. All hypotheses and measures were preregistered and can be accessed at https://osf.io/mpjkz/?view_only=7fd32ce53d434f4b8dbd0339579a8efa.

## 2. Material and methods

### 2.1 Participants

Ninety-two healthy participants (41 females, Age (M = 29.64, SD = 6.30, ranged from 18 to 47)) were recruited from the Los Angeles metropolitan area by the Caltech Conte Center for Social Decision-Making (P50 MH094258). All participants were fluent in English, had normal or corrected-to-normal vision and hearing, had Full Scale Intelligence Quotient greater than or equal to 90, had no first degree relative with schizophrenia or autism spectrum disorder, and had no history of developmental, psychiatric, or neurological disease. All participants provided written informed consent approved by the Institutional Review Board of the California Institute of Technology.

### 2.2 Magnetic Resonance Imaging

All MRI data was acquired using a 3T whole-body system (Magnetom TIM Trio, Siemens Medical Solutions, Malvern, PA) with a 32 channel receive head array at the Caltech Brain Imaging Center. Structural imaging data was acquired by the Imaging Core of the Caltech Conte Center for Social and Decision Neuroscience as part of a larger, multi-group consortium and analyzed retrospectively for this project. Structural images were acquired with one of two imaging protocols, corresponding to the first and second phases of the Caltech Conte Center (61 participants from Phase 1 and 31 participants from Phase 2). The Phase 1 protocol included two independent MP-RAGE acquisitions with TR/TE/TI = 1500/2.9/800 ms, flip angle = 10°, 1 mm isotropic voxels, 176 slab partitions, no in-plane GRAPPA, for a total imaging time of 12 minutes 52 seconds. The Phase 2 protocol included a single multi-echo MP-RAGE (MEMP-RAGE) acquisition with TR/TE/TI = 2530/1.6 to 7.2/1100 ms, flip angle = 7°, 0.9 mm isotropic voxels, 208 slab partitions, in-plane GRAPPA R = 2, for a total imaging time of 6 minutes 3 seconds. Both protocols generated T1-weighted structural images with comparable tissue contrast, SNR (following image or echo averaging) and voxel dimensions.

### 2.3 Estimation of cortical and subcortical volumes

Individual structural images were segmented and the cortical gray matter ribbon parcellated using the recon-all pipeline from Freesurfer v6.0.0 (Fischl, 2012). The pipeline initially registered and averaged the two separate T^1^-weighted images from the Phase 1 protocol prior to subsequent processing. Images from Phase 1 and Phase 2 protocols were processed independently and all images were resampled isotropically to 1 mm voxels prior to RF bias field correction and tissue segmentation. One hundred and forty-eight cortical gray matter parcel volumes (74 parcellations per hemisphere) corresponding to the Destrieux 2009 atlas (Destrieux, Fischl, Dale, & Halgren, 2010), seventeen subcortical region volumes, and estimated total intracranial volumes were compiled from the Freesurfer output for subsequent analysis in *R*. All cortical and subcortical volumes were normalized with respect to estimated total intracranial volume.

### 2.4 Social network index

The social network metric used in the present study is a subscale of the social network index, or SNI (Cohen, Doyle, Skoner, Rabin, & Gwaltney, 1997). This metric is a self-report questionnaire that quantifies the number of people participants saw or talked to at least once every two weeks in 12 different social relationships (e.g., spouse, children, relative, friend, neighbor, workmate). Participants from Phase 1 and Phase 2 did not differ in mean SNI (*t* = 0.93, *p* = 0.355; two-sample two-sided t-test). In addition to the SNI, we also asked participants to provide the modes of communication (e.g., face-to-face conversation, text, voice/video chat, social media) and types of support (e.g., emotional support, physical assistance, advice/information, companionship) used in those social relationships. Those variables were measured for the purpose of exploring whether SNI might be also associated with individual differences in modes of communication and types of support, as preregistered (see Appendix A).

### 2.5 Psychological measures

The cognitive ability of participants was measured with the Wechsler Abbreviated Scales of Intelligence-II (Wechsler, 2011), deriving two scores, verbal comprehension (M = 109.20, SD = 10.02) and perceptual reasoning (M = 104.80, SD = 10.86). The emotional intelligence (EI) of participants was measured with the Mayer-Salovey-Caruso Emotional Intelligence Test (Mayer, Salovey, & Caruso, 2002), deriving two sub-scores, experiential EI (M = 103.60, SD = 14.48) and strategic EI (M = 99.49, SD = 10.54). The empathy level of participants was measured with the Empathy Quotient (Baron-Cohen & Wheelwright, 2004) [M = 50.84, SD = 12.05]. The personality of participants was measured with the Sixteen Personality Factor Questionnaire (Cattell, Eber, & Tatsuoka, 1970; Russell, Karol, & Institute for Personality and Ability Testing, 2002), deriving five global scores, extraversion (M = 5.62, SD = 1.85), independence (M = 6.14, SD = 1.67), tough-mindedness (M = 4.35, SD = 1.60), self-control (M = 4.35, SD = 1.38), and anxiety (M = 5.65, SD = 1.85). The affect of participants was measured with the Positive and Negative Affect Schedule (Watson, Clark, & Carey, 1988), deriving two scores, positive affect (M = 31.68, SD = 8.43) and negative affect (M = 12.53, SD = 4.03). The stress level of participants was measured with the Perceived Stress Scale (Cohen, Kamarck, & Mermelstein, 1983) [M = 12.36, SD = 6.52]. The depression severity of participants was measured with the Beck Depression Inventory-II (Beck, Steer, & Brown, 1996) [M = 5.08, SD = 5.60]. The trait anxiety of participants was measured with the State-Trait Anxiety Inventory (Speilberger, Gorusch, Lushene, Vagg, & Jacobs, 1983) [M = 34.96, SD = 9.31].

### 2.6 Predictive modeling framework

To comprehensively understand the relationship between SNI and GMV, we carried out three independent analyses using three different predictive modeling frameworks (Figure 1). Framework 1 follows our pre-registered analysis plan and performed multivariate analysis (ridge regression) with cross-validation and feature selection. As recommended by recent research (Finn et al., 2015), we used univariate Pearson’s correlation between each feature and SNI as a criterion for feature selection. Specifically, we had an outer cross-validation loop that randomly split the data into training (80%) and test (20%) sets for 2000 iterations; in each outer loop iteration, the univariate Pearson’s correlation between each feature and SNI was assessed using the training data, and features that showed significant correlations with SNI (*p* < 0.05) were selected to construct a ridge regression model to predict SNI; the prediction accuracy of the model was then assessed using the test data. The hyperparameter (regularization penalty) of ridge regression was tuned using a nested cross-validation loop: the training data from the outer cross-validation loop were further randomly split into inner-training (80%) and inner-test (20%) for 20 iterations, and the optimal hyperparameter value was selected among 20 values in the interval of [1, 10000] across the 20 iterations.

**Fig. 1.**
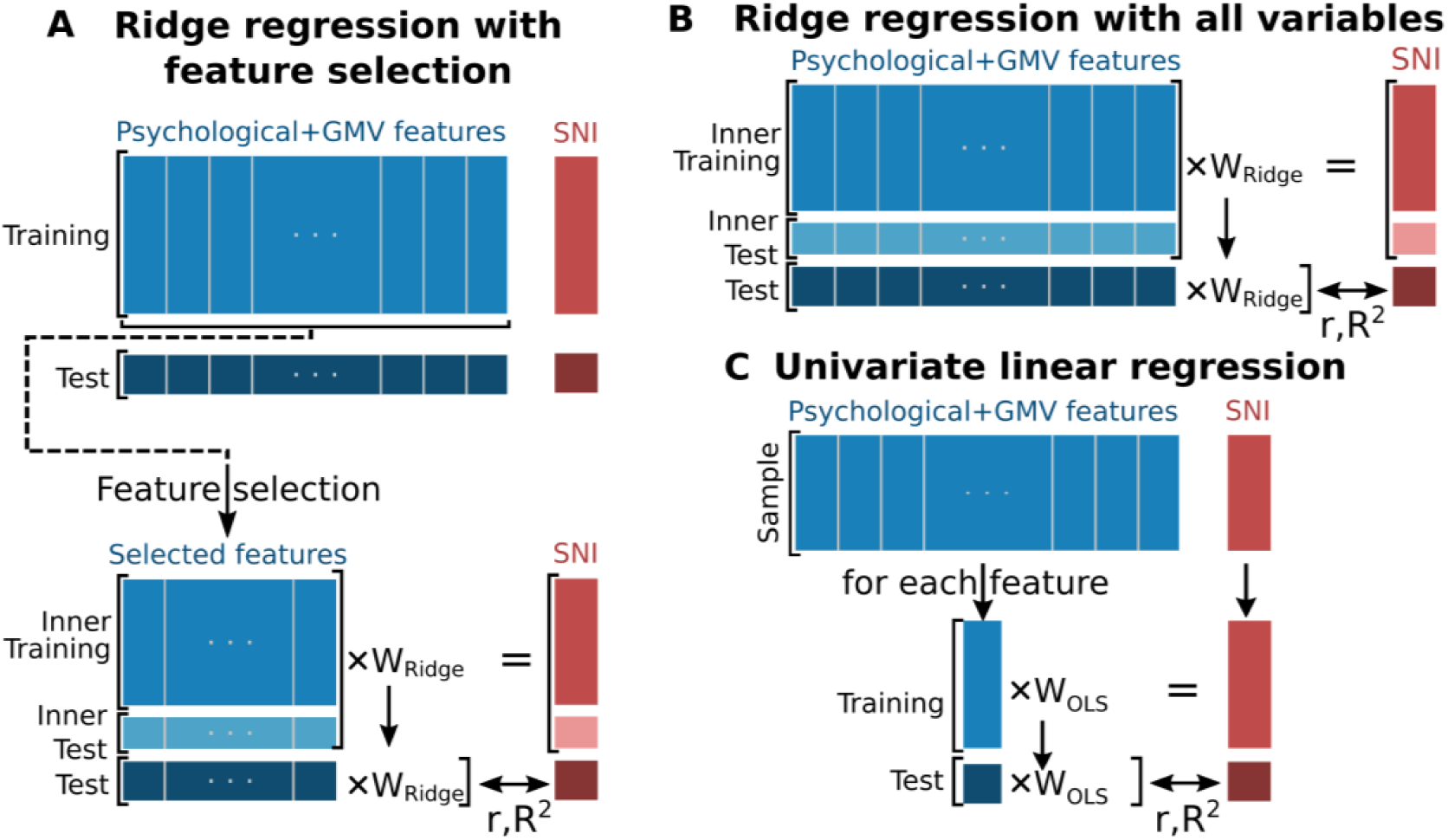
Illustration of three predictive modeling frameworks. (**A**) Framework 1 performed ridge regression with cross-validation using selected features. Features were selected within the cross-validation loop based on univariate correlations. The hyperparameter of ridge regression was tuned using a nested cross-validation loop. (**B**) Framework 2 performed ridge regression with cross-validation using all features. (**C**) Framework 3 performed univariate ordinary least-squares linear regression between each feature and SNI within the cross-validation loop.

To address the concern that the feature selection procedure might have omitted some features that did have associations with SNI, Framework 2 performed ridge regression with cross-validation without feature selection: the same procedures as in Framework 1 were used to construct the outer cross-validation loop and to tune the hyperparameter of ridge regression, except that the ridge regression model was fitted with all features in each iteration instead of selected features. To address the concern that the weights produced by multivariate models such as ridge regression could be misleading in the presence of correlated noise (Haufe et al., 2014; Kriegeskorte & Douglas, 2019), Framework 3 performed univariate linear regressions between every feature and SNI with cross-validation; cross-validation was constructed following the same procedures as in the first two frameworks for the outer cross-validation loop.

The prediction accuracy of each framework was assessed with two measures, Pearson’s *r* and prediction *R*^2^. Pearson’s *r* assessed the correlation between observed and predicted values of SNI in the test data. Prediction *R*^2^ measured the improvement of predicting SNI with our frameworks over the observed mean of SNI in the test data. The final reported prediction accuracy for each framework was averaged over the 2000 (outer loop) cross-validation splits. The p-values of prediction accuracies and model coefficients were calculated from permutations, where the null distributions were generated by randomly permuting the SNI labels across the sample for 10,000 iterations and in each iteration repeating all the analysis steps of a predictive framework. We handled multiple comparisons by correcting for false discovery rate (*q* < 0.05), which was applied when multiple features were tested for associations with SNI independently (i.e., univariate correlations in Framework 3) as well as when they were tested jointly (i.e., model coefficients in Frameworks 1 and 2). We handled the only binary feature, gender, by both removing the feature (which generated the results we reported here) and stratification (i.e., the training and test sets in cross-validation had approximately equal number of males and females); results from stratification corroborated those reported in the present paper. All analysis codes can be accessed at the Open Science Framework https://osf.io/zumwt/?view_only=4f11ca10ed5947c1be1ecdea57cfdff3.

## 3. Results

As preregistered, we first analyzed whether individual differences in SNI could be predicted by demographic characteristics and psychological measures alone. An exploratory factor analysis showed that a six-dimensional structure underlies the common variance of these eighteen psychological/demographic features (negative affect, cognitive control, extraversion, emotional intelligence, education, age and gender, see Appendix B). Analyses across all three frameworks consistently indicated that these eighteen psychological/demographic features alone did not predict SNI (see Appendix C).

Next, we inspected whether cortical and subcortical GMV together with psychological/ demographic features could predict individual differences in SNI. Analyses from Framework 3 showed that the effect size of every feature was weak, and none of the features alone predicted SNI after correcting for multiple comparisons (Table 2; see Appendix D for results of every feature). While univariate analyses generated model coefficients that were straightforward to interpret, they left open the question of whether multiple features combined might predict SNI. Analyses from Framework 1 and 2 showed that features in their entirety did not predict SNI either (Fig. 2).

**Table 2.**
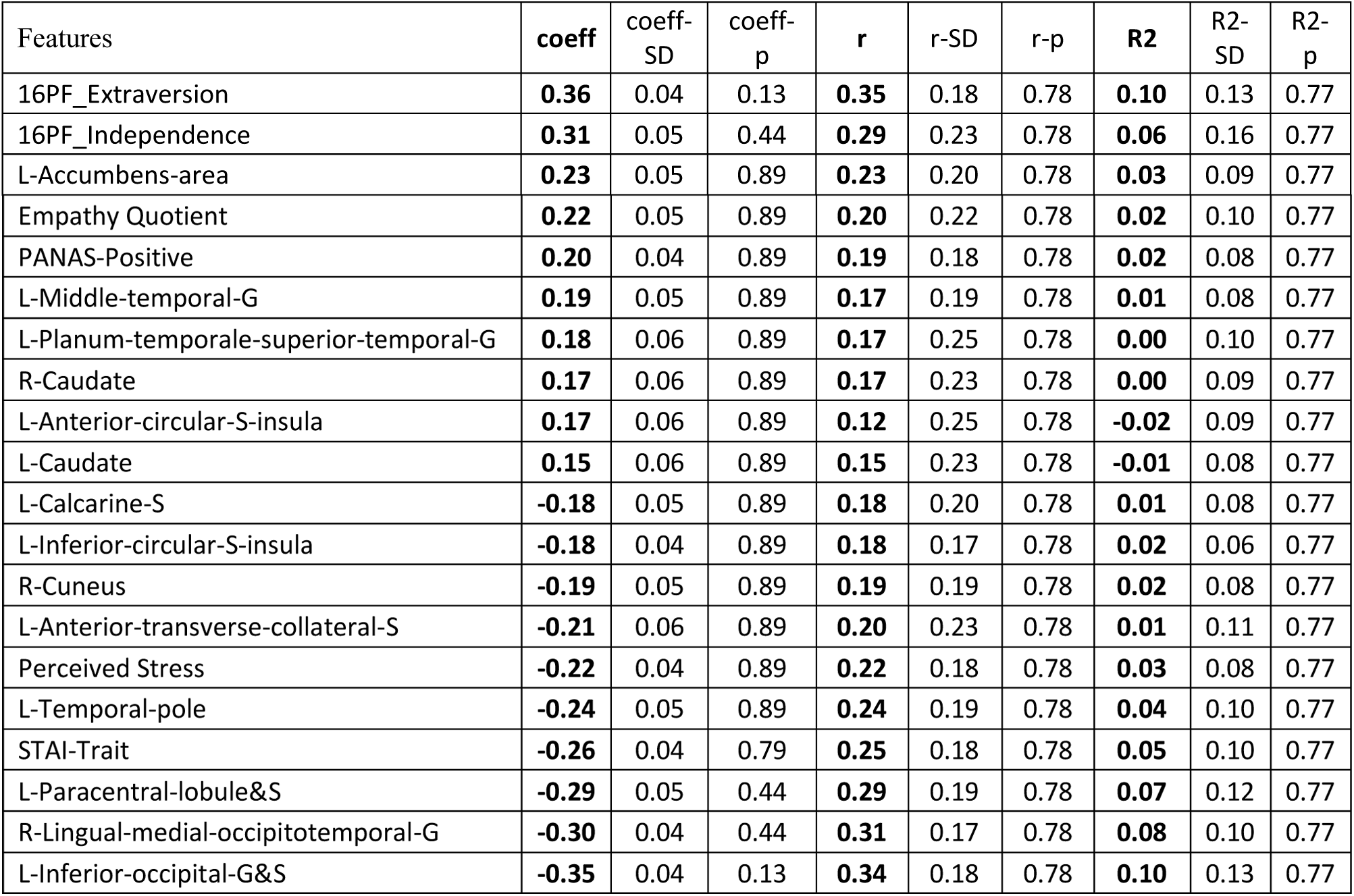
Results from univariate analyses of Framework 3. Model coefficients and prediction accuracies (with SDs, and p-values corrected for FDR) of the top ten features with the largest positive and negative effect sizes. Abbreviations: L left, R right, G gyrus/gyri, S sulcus/sulci, coeff coefficient.

**Fig. 2.**
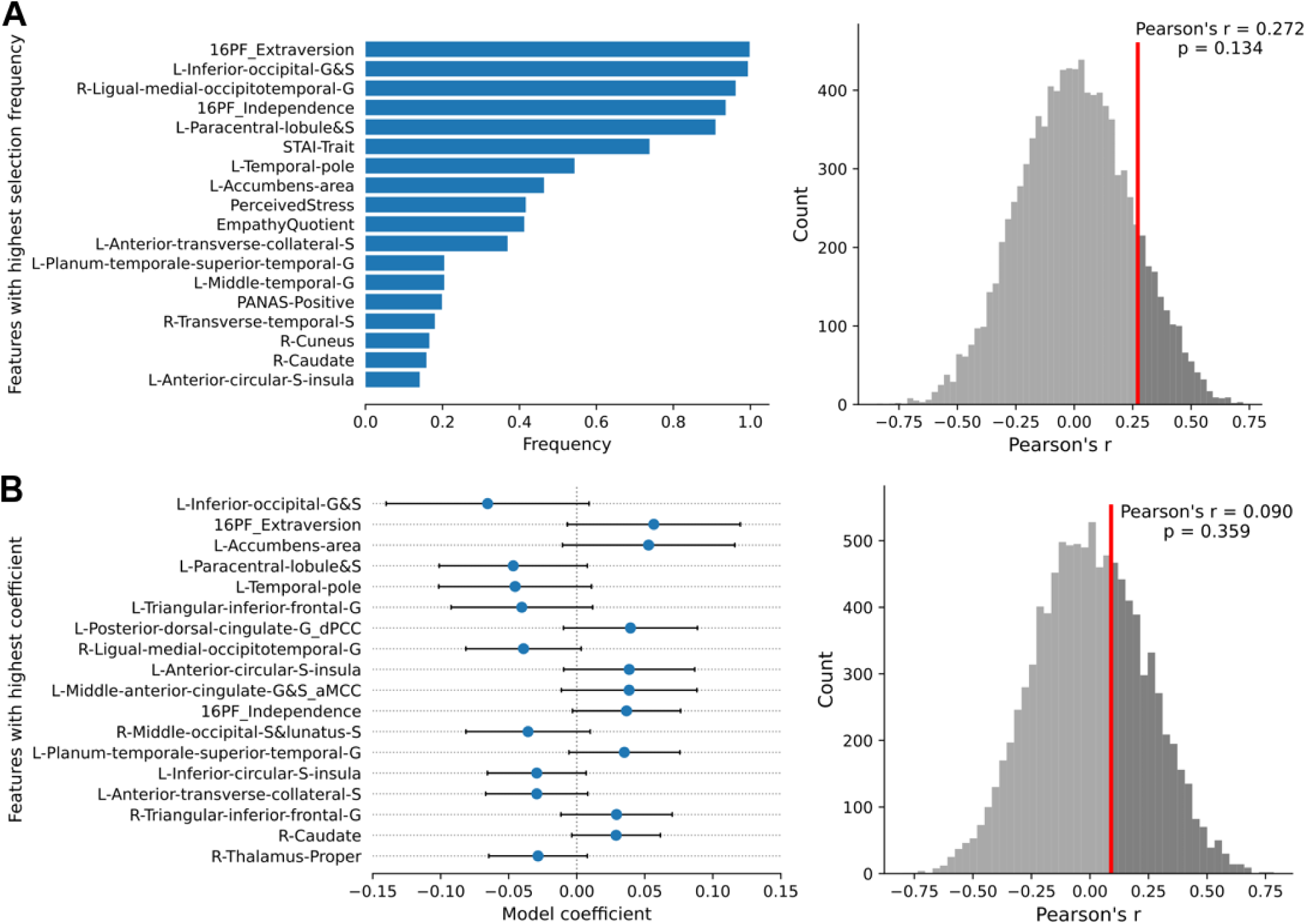
Predicting SNI with all GMV and psychological/demographic features. Results from analyses of Framework 1. The selection frequency (blue bars) of the top (most frequently selected) eighteen features over the 2000 iterations of the outer cross-validation loop (left) and the mean prediction accuracy (red vertical line, assessed with Pearson’s r) averaged over the 2000 outer cross-validation iterations compared to the null distribution generated with permutation (right). The mean prediction accuracy assessed with prediction *R*^2^ = 0.060, p = 0.136. (**B**) Results from analyses of Framework 2. Model coefficients (blue dots) and standard deviations (black bars) of the top eighteen features (left) and the mean prediction accuracy (red vertical line, assessed with Pearson’s r) averaged over the 2000 outer cross-validation iterations compared to the null distribution generated with permutation (right). The mean prediction accuracy assessed with prediction *R*^2^ = -0.023, p = 0.404.

While our study used a predictive framework (using cross-validation), we also recognize the value of descriptive effect sizes in providing results that could be used to formulate hypotheses to be tested in future studies. To that end, we also show, for every cortical and subcortical region over the brain, the univariate effect size of the correlation between SNI and GMV estimated using all data (Figure 3, Appendix E).

**Fig. 3.**
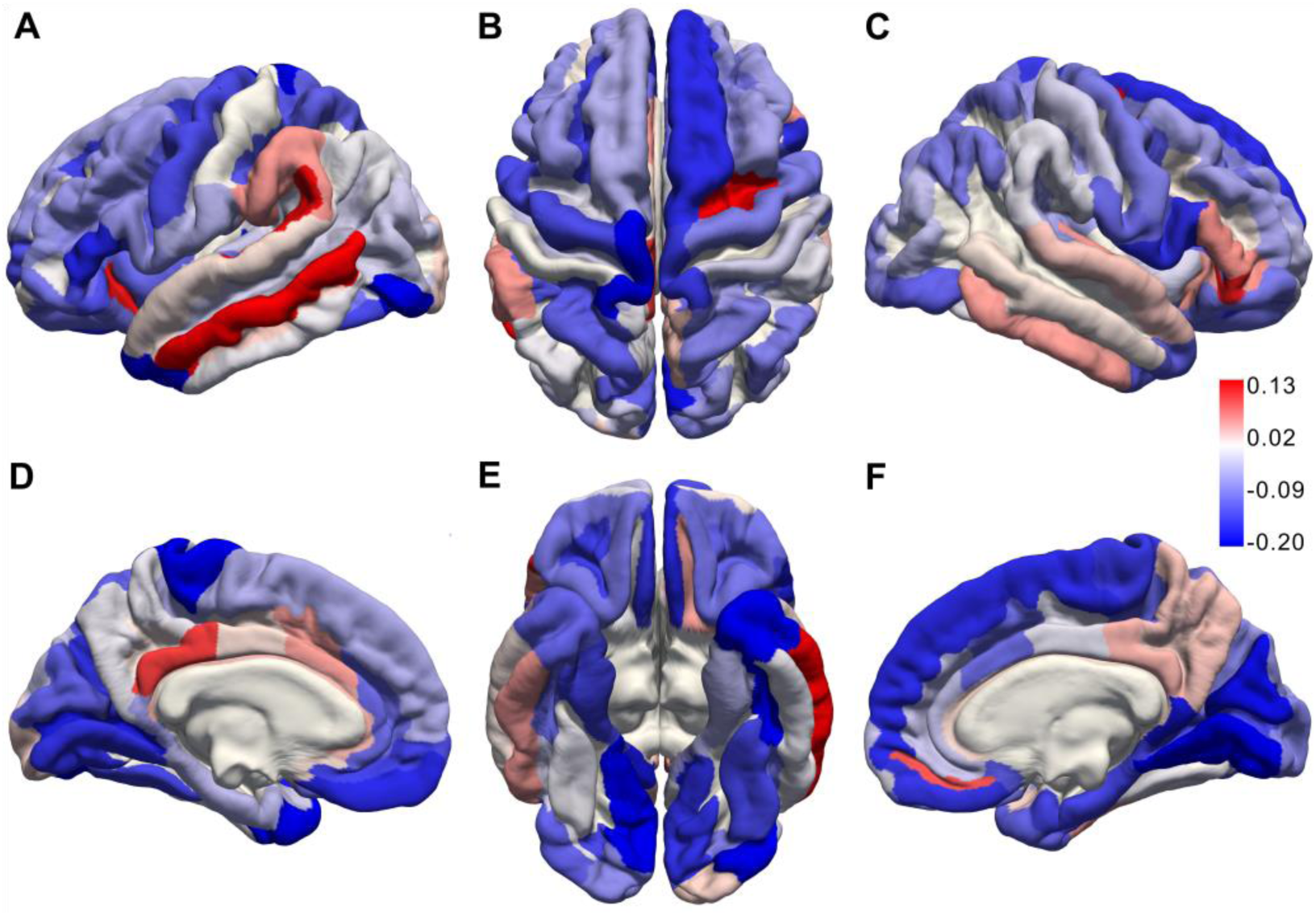
Descriptive effect sizes between SNI and every cortical GMV. The descriptive effect size of the univariate associations between all cortical regions and SNI are shown to provide background for future studies that could test hypotheses based on these results. Four renderings of the univariate Pearson correlations (uncorrected) between individual cortical regions and SNI are projected on the pial surface for (**A**) the lateral view of the left hemisphere, (**B**) the superior view of both hemispheres, (**C**) the lateral view of the right hemisphere, (**D**) the medial view of the left hemisphere, (**E**) the inferior view of both hemispheres, and (**F**) the medial view of the right hemisphere. These effect sizes provide recommendations for the sample sizes required to test associations between specific cortical regions and SNI, shown in Appendix E.

## 4. Discussion

Following our preregistration, we applied a data-driven approach to comprehensively examine the relationship between SNI and demographic, psychological, cortical and subcortical GMV features, using three different predictive modeling frameworks (Fig. 1). In our sample of healthy adult humans, no evidence was found that any feature was significantly associated with SNI after multiple comparison corrections (Fig. 2 and Table 2). It is important to note that whether a given effect will be detected as significant or not is of course highly dependent on the sample size (i.e., the larger the sample size, the easier it is to detect a given effect size); similarly, estimated effect sizes and their statistical significance will vary depending on the analysis frameworks (e.g., methods for model construction and multiple comparison corrections). Our study used a comparatively large sample, tested three different predictive modeling frameworks, and included pre-registration to verify the degrees of freedom in our analyses and to facilitate sharing of data and codes. Regardless of statistical significance, we note that the estimated effect size of most features, in particular 159 of the 165 cortical and subcortical GMV features, were very weak, even when assessed with the simplest univariate correlation method (absolute values less than 0.20; see Fig. 3 and Appendix E). These findings do not demonstrate that there is no association between GMV and SNI, but they do urge caution in interpreting prior reports of such associations. We suggest that additional studies are needed on this topic, and that a future meta-analysis based on all studies will be required to obtain a more accurate estimate of the true effect sizes on this topic.

Three features reported in previous studies (Table 1; Asendorpf & Wilpers, 1998) to have a significant positive association with social network metrics—extraversion, left middle temporal gyrus GMV, and left anterior insula GMV—and one feature reported in previous studies (Nabi, Prestin, & So, 2013) to have a significant negative association with social network metrics— perceived stress—indeed showed relatively larger effect sizes in expected directions among the features in our sample (Table 2). However, those effect sizes were still very weak and were not significant in our study after multiple comparison corrections. The left temporal pole GMV has also been reported to positively correlate with social network metrics (Table 1); though this region showed a relatively larger effect size among our features (Table 2), it was in the opposite direction from what has been reported previously (negative). Previously unreported regions in the left occipital cortex also showed a relatively larger negative effect among the features. We do not have an explanation for these negative effects and suggest that they may well be statistically unreliable effects that turned up by chance given that we sampled all brain regions—indeed, these negative effects were not significant after multiple comparison corrections. Nonetheless, the specific GMV regions discussed in this section should serve as predictors in future hypothesis-driven studies that could focus on one or several of these features.

We previously noted the reliable positive correlation between neocortex volume and social group size found across species (Dunbar, 1998; Dunbar & Shultz, 2007), and that this finding might suggest the possibility that such a relationship would also exist across individuals within a single species such as humans. However, any reliable relationship between social network metrics for a specific individual and GMV is less plausible once we consider that social network metrics such as SNI in individual humans is quite changeable, fluctuating as people move to new locations, get a new job, or encounter other common transitions in their lives. Our failure to replicate previously reported effects of GMV fit with this picture, and raise the possibility that many prior findings might be false positives. Measures other than the SNI that could obtain more temporally stable metrics related to social network size would seem better suited for investigating associations with GMV. Alternatively, more dynamic measures of brain function, rather than structure, would seem better suited for exploring associations with SNI. We would expect that functional measures (or possibly others, such as from diffusion MRI) might well yield associations with SNI (Bickart, Hollenbeck, Barrett, & Dickerson, 2012; Dziura & Thompson, 2014; Hampton, Unger, Von Der Heide, & Olson, 2016; Pillemer, Holtzer, & Blumen, 2017).

The non-significant effects of many previously reported regions that we found in the present study might be related to several limitations of our study, and of course do not demonstrate that there is no effect. First, compared to the seminal study that reported a correlation between amygdala volume and SNI (Bickart et al., 2011), our sample has a narrower age range, which might result in less variability in amygdala volume and therefore lower power to detect an association between amygdala volume and SNI. Second, all cortical and subcortical GMV used in the present study were measured based on automated segmentations from FreeSurfer without any manual correction (although we did carry out manual checks on a subset of the segmentation results to verify their quality). This procedure has been shown to be no less accurate than manual labeling (Bickart et al., 2011; Fischl et al., 2002), yet potential errors in segmentation might have also reduced power to find a relationship between SNI and GMV.

We conclude with three recommendations for future research. First, studies attempting to test the relationship between social network metrics and structural brain measures in humans should first ensure that their respective sets of measures are approximately matched in terms of temporal stability (e.g., using structural MRI predictors for temporally stable network measures, but functional MRI predictors for metrics such as the SNI). Second, given concerns about false positives when testing for associations between multiple regions and social network metrics, future studies should try to preregister their hypotheses—and in particular, methods of correcting for multiple comparisons—before conducting the analyses (Nosek, Ebersole, DeHaven, & Mellor, 2018). Such preregistered studies, if focused on specific neuroanatomical regions, should include sample sizes sufficiently large to detect the hypothesized associations (Appendix E). As well, it is essential for studies to share all data and codes (e.g., through OSF) so that future meta-analyses can capitalize on all accumulated findings. Third, future studies should focus on understanding the mechanisms that might explain any association between social network metrics and GMV of some regions in the brain. For example, some studies have suggested that mentalizing might mediate such associations (Powell, Lewis, Roberts, García-Fiñana, & Dunbar, 2012). This hypothesis could be tested with a more formal structural equation model, namely, that GMV in brain regions thought to subserve mentalizing causes individual differences in actual mentalizing ability in real life, which in turn has a causal effect on how many people an individual associates with in social networks. Future studies employing longitudinal designs (e.g., repeatedly measuring social network metrics and GMV over years), mediation analyses, and meta-analyses would shed new light on the mechanisms underlying the relationship between social network metrics and structural brain measures.

## Appendices

## Appendix A. Correlations between SNI, modes of communication, and types of support

As preregistered, we explored the relationship between SNI and modes of communication and types of support in the 12 social relationships. We collected these measures in two independent samples of participants (an in-lab sample with 57 participants and an online-sample with 101 participants), reporting findings in both samples as replications. Besides the Social Network Index (from which we derived all three scores: the number of people in network, network diversity, and the number of embedded networks), participants were asked whether they used any of the seven modes of communication (face-to-face conversation, text, voice/video chat, email, social media, gaming, touch) in each social relationship, and furthermore whether they received or provided any of the five types of support (emotional support, physical/material assistance, advice/information, appraisal, companionship) in each social relationship. A summary score for each mode and each type of support was derived by averaging the responses across all social relationships. Numbers indicate the average correlation across the two samples. Numbers were colored only if the correlations were significant in both samples.

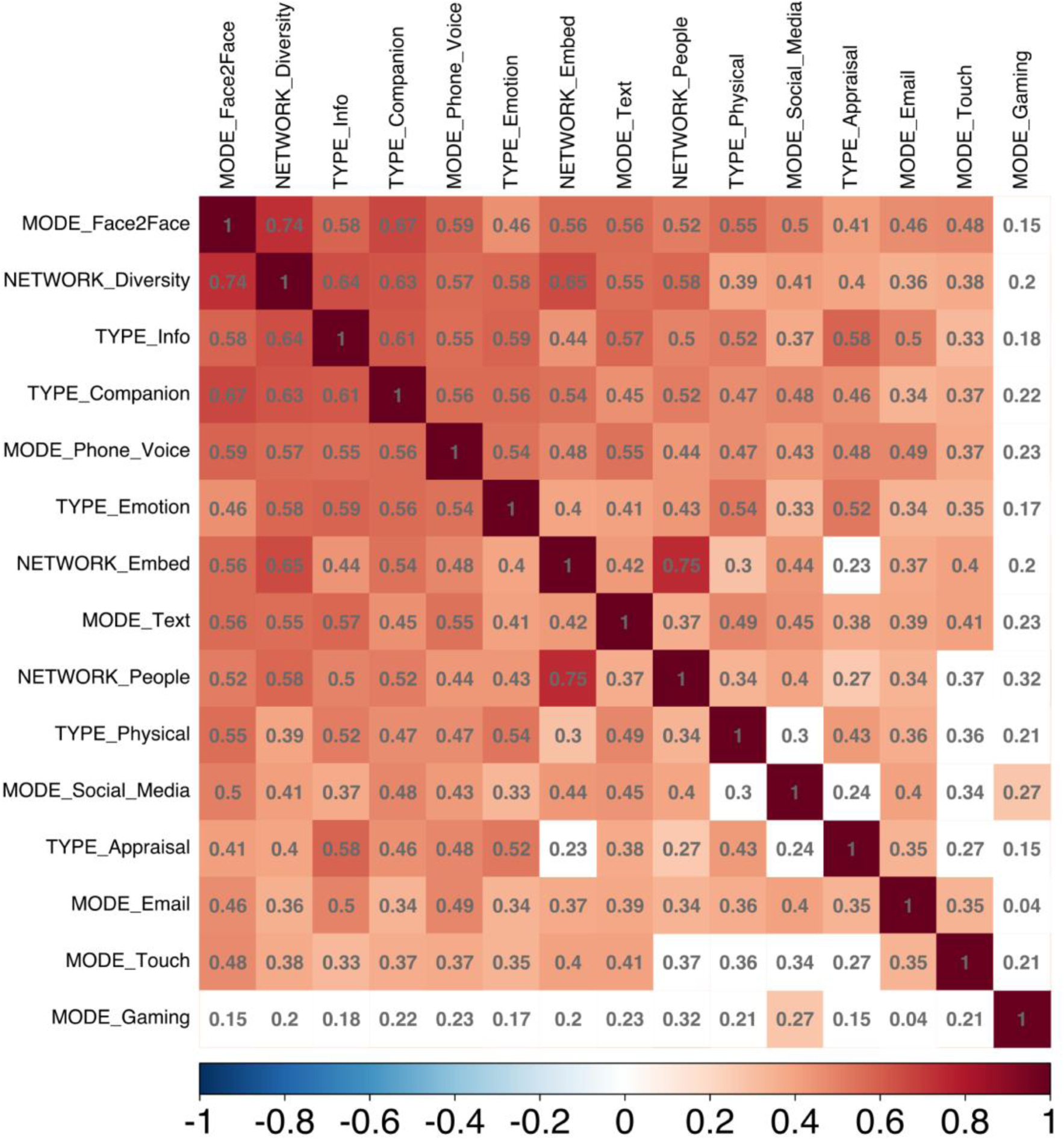

## Appendix B. Exploratory factor analysis on demographic characteristics and psychological measures

Cattell’s scree test and Kaiser’s rule both indicated that a six-factor structure underlies the common variance in the data. Therefore, we applied exploratory factor analysis to extract six factors using the minimal residual method. The solutions were rotated with oblimin for interpretability. Each column plotted the strength of the factor loadings (x-axis, absolute value) across all demographic characteristics and psychological measures. The color of the bar indicated the sign of the loading (red for positive and blue for negative; more saturated for higher absolute values).

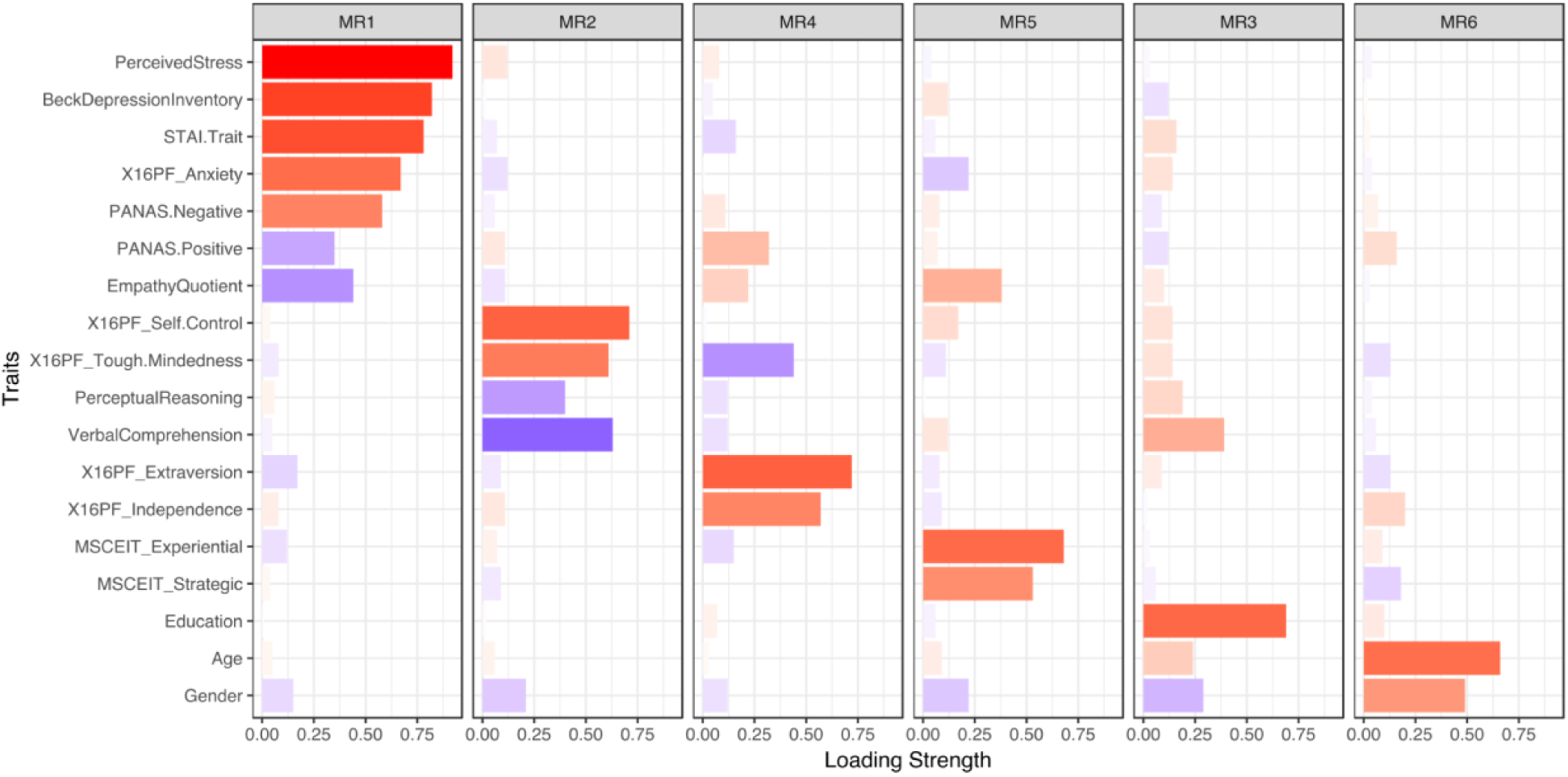

## Appendix C. Predicting SNI with demographic and psychological features alone

(**A**) The selection frequency of each feature (left) and the model prediction accuracy compared with the null distribution (right) obtained from Framework 1. The model accuracy assessed with prediction *R*^2^ = 0.085, *p* = 0.072. (**B**) The model coefficients and standard deviations (left) and the model prediction accuracy compared with the null distribution (right) obtained from Framework 2. The model accuracy assessed with prediction *R*^2^ = 0.054, *p* = 0.185. (**C**) The model coefficients and accuracies (assessed with both Pearson’s *r* and prediction *R*^2^) with SDs and p-values corrected for FDR obtained from Framework 3.

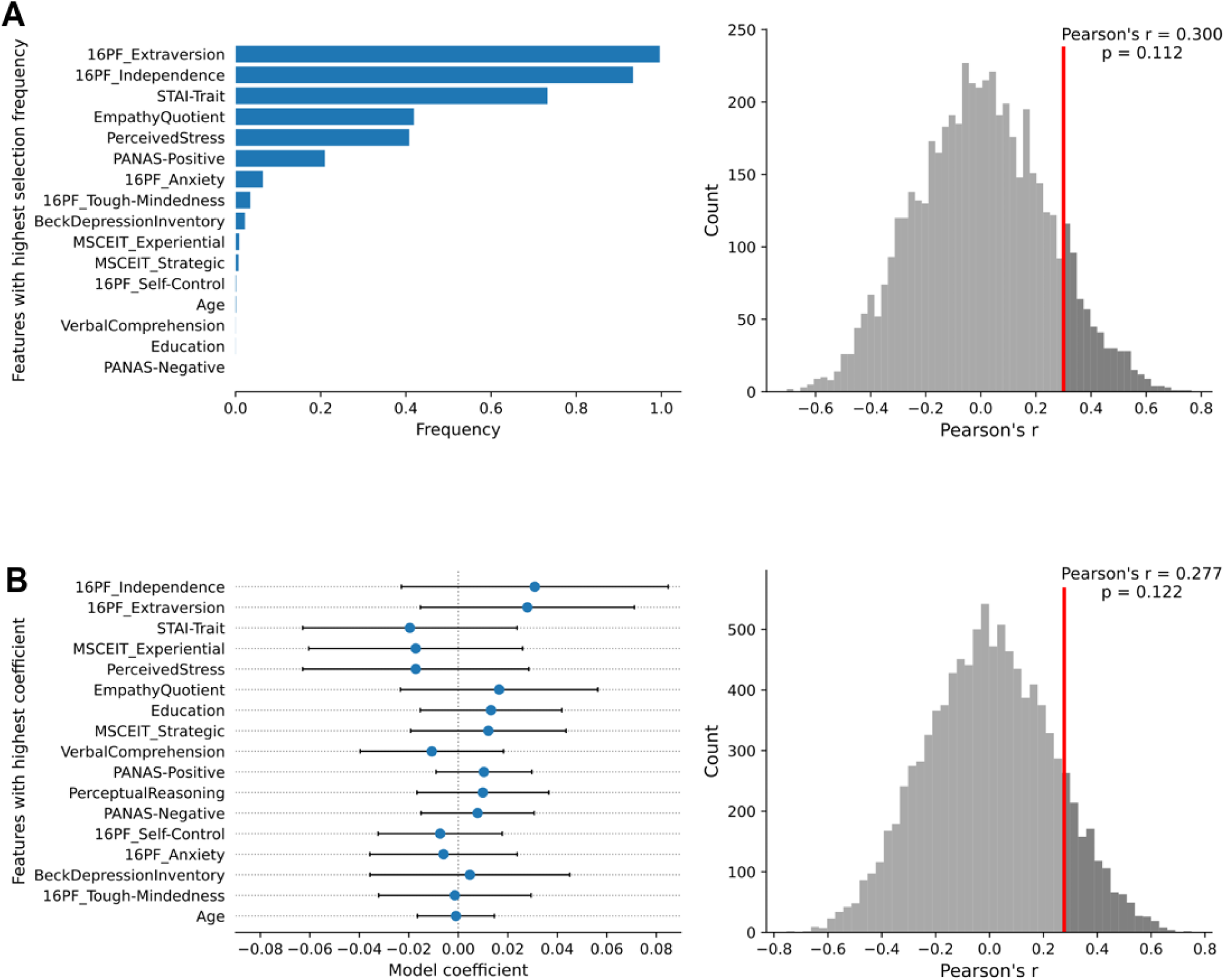
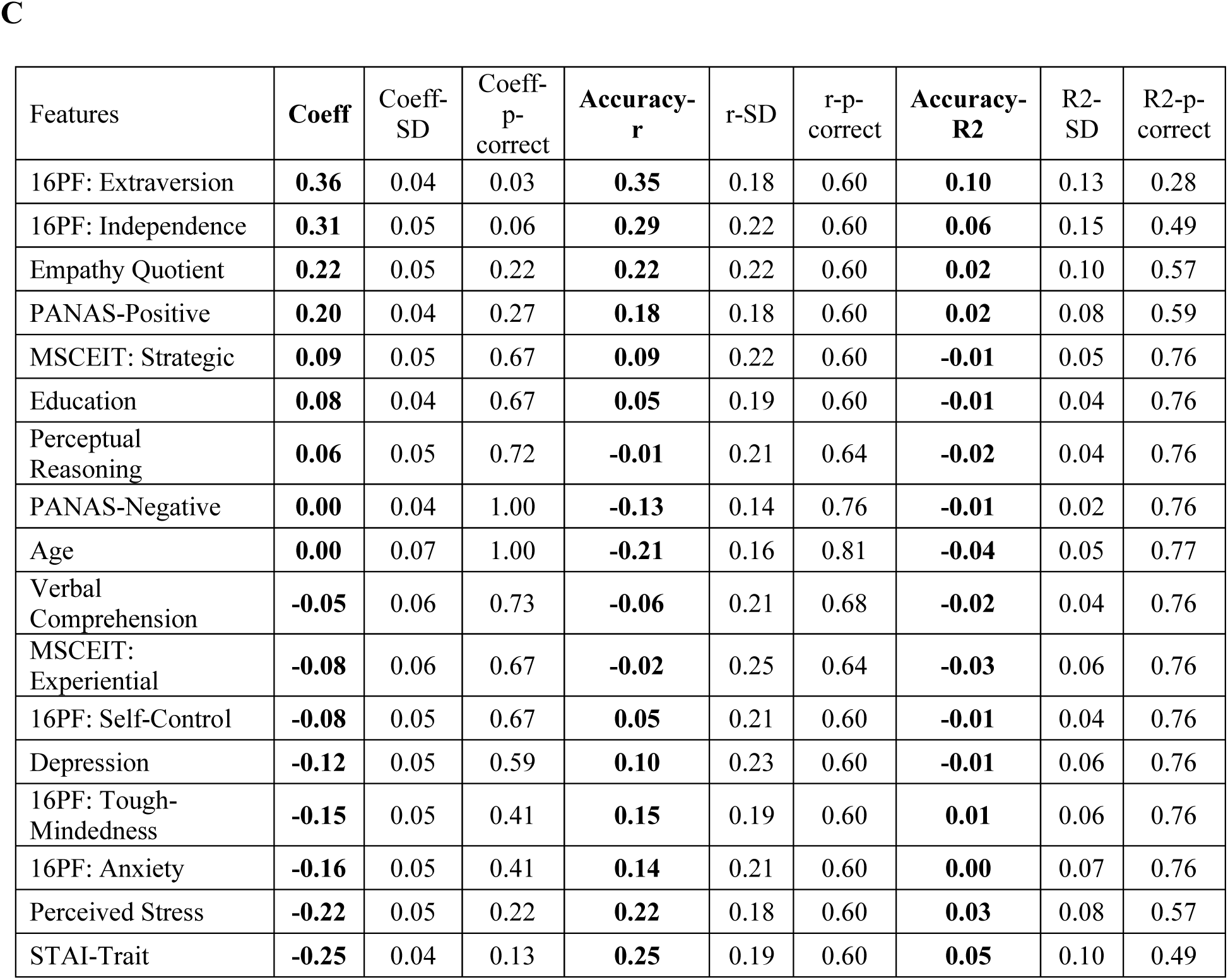

## Appendix D. Predicting SNI with GMV and all other features using Framework 3

The model coefficients and accuracies (assessed with both Pearson’s *r* and prediction *R*^2^) with SDs and p-values corrected for FDR obtained from Framework 3.

| Features | coeff | coeff-SD | coeff-p | r | r-SD | r-p | R2 | R2-SD | R2-p |
| --- | --- | --- | --- | --- | --- | --- | --- | --- | --- |
| 16PF_Extraversion | **0.36** | 0.04 | 0.13 | **0.35** | 0.18 | 0.78 | **0.10** | 0.13 | 0.77 |
| 16PF_Independence | **0.31** | 0.05 | 0.44 | **0.29** | 0.23 | 0.78 | **0.06** | 0.16 | 0.77 |
| L-Accumbens-area | **0.23** | 0.05 | 0.89 | **0.23** | 0.20 | 0.78 | **0.03** | 0.09 | 0.77 |
| Empathy Quotient | **0.22** | 0.05 | 0.89 | **0.20** | 0.22 | 0.78 | **0.02** | 0.10 | 0.77 |
| PANAS-Positive | **0.20** | 0.04 | 0.89 | **0.19** | 0.18 | 0.78 | **0.02** | 0.08 | 0.77 |
| L-Middle-temporal-G | **0.19** | 0.05 | 0.89 | **0.17** | 0.19 | 0.78 | **0.01** | 0.08 | 0.77 |
| L-Planum-temporale-superior-temporal-G | **0.18** | 0.06 | 0.89 | **0.17** | 0.25 | 0.78 | **0.00** | 0.10 | 0.77 |
| R-Caudate | **0.17** | 0.06 | 0.89 | **0.17** | 0.23 | 0.78 | **0.00** | 0.09 | 0.77 |
| L-Anterior-circular-S-insula | **0.17** | 0.06 | 0.89 | **0.12** | 0.25 | 0.78 | **-0.02** | 0.09 | 0.77 |
| L-Caudate | **0.15** | 0.06 | 0.89 | **0.15** | 0.23 | 0.78 | **-0.01** | 0.08 | 0.77 |
| L-Posterior-dorsal-cingulate-G_dPCC | **0.14** | 0.06 | 0.89 | **0.10** | 0.23 | 0.78 | **-0.01** | 0.07 | 0.77 |
| R-Superior-precentral-S | **0.13** | 0.06 | 0.89 | **0.07** | 0.24 | 0.78 | **-0.02** | 0.07 | 0.77 |
| L-Amygdala | **0.12** | 0.06 | 0.89 | **0.10** | 0.24 | 0.78 | **-0.02** | 0.07 | 0.77 |
| R-Orbital-inferior-frontal-G | **0.11** | 0.05 | 0.89 | **0.10** | 0.20 | 0.78 | **-0.01** | 0.05 | 0.77 |
| R-Pallidum | **0.11** | 0.05 | 0.89 | **0.10** | 0.21 | 0.78 | **-0.01** | 0.06 | 0.77 |
| MSCEIT_Strategic | **0.09** | 0.05 | 0.92 | **0.07** | 0.23 | 0.78 | **-0.01** | 0.05 | 0.77 |
| R-Amygdala | **0.09** | 0.06 | 0.93 | **0.03** | 0.24 | 0.78 | **-0.02** | 0.06 | 0.77 |
| R-Suborbital-S | **0.08** | 0.05 | 0.96 | **0.02** | 0.22 | 0.78 | **-0.02** | 0.05 | 0.77 |
| Education | **0.08** | 0.04 | 0.96 | **0.05** | 0.19 | 0.78 | **-0.01** | 0.03 | 0.77 |
| L-Supramarginal-G | **0.07** | 0.06 | 0.96 | **-0.02** | 0.24 | 0.78 | **-0.02** | 0.05 | 0.77 |
| L-Middle-anterior-cingulate-G&S_aMCC | **0.07** | 0.04 | 0.97 | **0.02** | 0.20 | 0.78 | **-0.01** | 0.03 | 0.77 |
| Perceptual Reasoning | **0.06** | 0.05 | 0.97 | **-0.02** | 0.21 | 0.78 | **-0.02** | 0.04 | 0.77 |
| L-Medial-orbital-S | **0.06** | 0.05 | 0.97 | **-0.02** | 0.21 | 0.78 | **-0.02** | 0.04 | 0.77 |
| R-Triangular-inferior-frontal-G | **0.05** | 0.05 | 0.98 | **-0.03** | 0.19 | 0.78 | **-0.02** | 0.03 | 0.77 |
| R-Lateral-orbital-S | **0.05** | 0.05 | 0.98 | **-0.04** | 0.19 | 0.78 | **-0.02** | 0.03 | 0.77 |
| R-Anterior-transverse-temporal-G | **0.05** | 0.05 | 0.98 | **-0.06** | 0.19 | 0.78 | **-0.02** | 0.03 | 0.77 |
| L-Pericallosal-S | **0.05** | 0.05 | 0.98 | **-0.07** | 0.20 | 0.78 | **-0.02** | 0.03 | 0.77 |
| L-Pallidum | **0.04** | 0.06 | 0.98 | **-0.10** | 0.21 | 0.78 | **-0.03** | 0.04 | 0.77 |
| R-Inferior-temporal-G | **0.04** | 0.06 | 0.98 | **-0.10** | 0.19 | 0.78 | **-0.03** | 0.04 | 0.77 |
| R-Anterior-circular-S-insula | **0.04** | 0.06 | 0.98 | **-0.10** | 0.21 | 0.78 | **-0.02** | 0.04 | 0.77 |
| R-Inferior-temporal-S | **0.04** | 0.05 | 0.98 | **-0.07** | 0.19 | 0.78 | **-0.02** | 0.03 | 0.77 |
| R-Hippocampus | **0.04** | 0.05 | 0.98 | **-0.10** | 0.19 | 0.78 | **-0.02** | 0.03 | 0.77 |
| R-Posterior-dorsal-cingulate-G_dPCC | **0.03** | 0.04 | 0.98 | **-0.08** | 0.16 | 0.78 | **-0.02** | 0.03 | 0.77 |
| L-Inferior-temporal-S | **0.03** | 0.05 | 0.98 | **-0.10** | 0.17 | 0.78 | **-0.02** | 0.03 | 0.77 |
| L-Occipital-pole | **0.03** | 0.06 | 0.98 | **-0.16** | 0.19 | 0.78 | **-0.03** | 0.05 | 0.77 |
| R-Precuneus | **0.03** | 0.05 | 0.98 | **-0.14** | 0.16 | 0.78 | **-0.02** | 0.03 | 0.77 |
| R-Lateral-superior-temporal-G | **0.03** | 0.05 | 0.98 | **-0.12** | 0.17 | 0.78 | **-0.02** | 0.03 | 0.77 |
| R-Subparietal-S | **0.02** | 0.05 | 0.98 | **-0.13** | 0.15 | 0.78 | **-0.02** | 0.03 | 0.77 |
| L-Middle-posterior-cingulate-G&S_pMCC | **0.02** | 0.05 | 0.98 | **-0.14** | 0.15 | 0.78 | **-0.02** | 0.03 | 0.77 |
| R-Planum-polare-superior-temporal-G | **0.02** | 0.05 | 0.98 | **-0.13** | 0.14 | 0.78 | **-0.02** | 0.02 | 0.77 |
| L-Subcallosal-G | **0.02** | 0.05 | 0.98 | **-0.13** | 0.14 | 0.78 | **-0.02** | 0.02 | 0.77 |
| L-Lateral-superior-temporal-G | **0.02** | 0.05 | 0.98 | **-0.14** | 0.14 | 0.78 | **-0.02** | 0.03 | 0.77 |
| R-Pericallosal-S | **0.02** | 0.05 | 0.98 | **-0.15** | 0.15 | 0.78 | **-0.02** | 0.03 | 0.77 |
| L-Subparietal-S | **0.01** | 0.06 | 0.98 | **-0.17** | 0.16 | 0.78 | **-0.03** | 0.04 | 0.77 |
| L-Frontomarginal-G&S | **0.01** | 0.06 | 0.99 | **-0.19** | 0.15 | 0.79 | **-0.03** | 0.04 | 0.77 |
| R-Middle-temporal-G | **0.01** | 0.04 | 0.98 | **-0.12** | 0.12 | 0.78 | **-0.01** | 0.02 | 0.77 |
| R-Inferior-frontal-S | **0.01** | 0.05 | 0.99 | **-0.17** | 0.14 | 0.78 | **-0.02** | 0.03 | 0.77 |
| L-Middle-frontal-S | **0.01** | 0.06 | 0.99 | **-0.17** | 0.14 | 0.78 | **-0.03** | 0.03 | 0.77 |
| L-S-intermedius-primus | **0.00** | 0.06 | 1.00 | **-0.17** | 0.14 | 0.78 | **-0.03** | 0.03 | 0.77 |
| R-Anterior-occipital-S&preoccipital-notch | **0.00** | 0.05 | 1.00 | **-0.14** | 0.12 | 0.78 | **-0.02** | 0.03 | 0.77 |
| R-Central-S | **0.00** | 0.06 | 1.00 | **-0.18** | 0.14 | 0.78 | **-0.03** | 0.04 | 0.77 |
| R-Superior-temporal-S | **0.00** | 0.06 | 1.00 | **-0.18** | 0.14 | 0.78 | **-0.03** | 0.04 | 0.77 |
| Age | **0.00** | 0.07 | 1.00 | **-0.21** | 0.16 | 0.81 | **-0.04** | 0.05 | 0.78 |
| PANAS-Negative | **0.00** | 0.04 | 1.00 | **-0.14** | 0.13 | 0.78 | **-0.01** | 0.02 | 0.77 |
| L-Postcentral-G | **0.00** | 0.06 | 1.00 | **-0.17** | 0.14 | 0.78 | **-0.03** | 0.03 | 0.77 |
| L-Middle-occipital-S&lunatus-S | **-0.01** | 0.05 | 1.00 | **-0.15** | 0.13 | 0.78 | **-0.02** | 0.03 | 0.77 |
| L-Lateral-orbital-S | **-0.01** | 0.05 | 1.00 | **-0.15** | 0.14 | 0.78 | **-0.02** | 0.03 | 0.77 |
| L-Putamen | **-0.01** | 0.06 | 0.98 | **-0.17** | 0.15 | 0.78 | **-0.03** | 0.04 | 0.77 |
| R-Medial-orbital-S | **-0.01** | 0.05 | 0.98 | **-0.15** | 0.14 | 0.78 | **-0.02** | 0.03 | 0.77 |
| R-Lateral-occipitotemporal-G | **-0.01** | 0.06 | 0.98 | **-0.17** | 0.15 | 0.78 | **-0.03** | 0.04 | 0.77 |
| L-Inferior-temporal-G | **-0.02** | 0.05 | 0.98 | **-0.15** | 0.15 | 0.78 | **-0.02** | 0.03 | 0.77 |
| L-Long-insular-G&central-insula-S | **-0.02** | 0.05 | 0.98 | **-0.14** | 0.14 | 0.78 | **-0.02** | 0.03 | 0.77 |
| L-Precuneus | **-0.02** | 0.06 | 0.98 | **-0.16** | 0.16 | 0.78 | **-0.03** | 0.04 | 0.77 |
| L-Angular-G | **-0.02** | 0.06 | 0.98 | **-0.17** | 0.16 | 0.78 | **-0.03** | 0.04 | 0.77 |
| R-Frontomarginal-G&S | **-0.02** | 0.06 | 0.98 | **-0.17** | 0.16 | 0.78 | **-0.03** | 0.04 | 0.77 |
| R-Supramarginal-G | **-0.02** | 0.05 | 0.98 | **-0.13** | 0.15 | 0.78 | **-0.02** | 0.03 | 0.77 |
| R-Middle-posterior-cingulate-G&S_pMCC | **-0.02** | 0.06 | 0.98 | **-0.15** | 0.17 | 0.78 | **-0.03** | 0.04 | 0.77 |
| R-S-intermedius-primus | **-0.02** | 0.06 | 0.98 | **-0.15** | 0.16 | 0.78 | **-0.03** | 0.04 | 0.77 |
| L-Posterior-lateral-S | **-0.02** | 0.07 | 0.98 | **-0.18** | 0.17 | 0.78 | **-0.03** | 0.05 | 0.77 |
| L-Superior-precentral-S | **-0.02** | 0.04 | 0.98 | **-0.10** | 0.15 | 0.78 | **-0.01** | 0.02 | 0.77 |
| R-Superior-occipital-S&transverse-occipital-S | **-0.02** | 0.04 | 0.98 | **-0.11** | 0.15 | 0.78 | **-0.02** | 0.03 | 0.77 |
| R-Short-insular-G | **-0.03** | 0.06 | 0.98 | **-0.15** | 0.17 | 0.78 | **-0.03** | 0.04 | 0.77 |
| R-Lateral-occipitotemporal-S | **-0.03** | 0.05 | 0.98 | **-0.10** | 0.16 | 0.78 | **-0.02** | 0.03 | 0.77 |
| L-Anterior-occipital-S&preoccipital-notch | **-0.03** | 0.04 | 0.98 | **-0.09** | 0.16 | 0.78 | **-0.02** | 0.02 | 0.77 |
| L-Superior-temporal-S | **-0.03** | 0.06 | 0.98 | **-0.13** | 0.18 | 0.78 | **-0.03** | 0.04 | 0.77 |
| L-Planum-polare-superior-temporal-G | **-0.04** | 0.05 | 0.98 | **-0.10** | 0.18 | 0.78 | **-0.02** | 0.03 | 0.77 |
| R-Postcentral-G | **-0.04** | 0.06 | 0.98 | **-0.13** | 0.19 | 0.78 | **-0.03** | 0.04 | 0.77 |
| L-Superior-occipital-G | **-0.04** | 0.05 | 0.98 | **-0.09** | 0.18 | 0.78 | **-0.02** | 0.03 | 0.77 |
| L-Middle-occipital-G | **-0.04** | 0.05 | 0.98 | **-0.10** | 0.19 | 0.78 | **-0.02** | 0.04 | 0.77 |
| L-Anterior-transverse-temporal-G | **-0.04** | 0.05 | 0.98 | **-0.09** | 0.18 | 0.78 | **-0.02** | 0.03 | 0.77 |
| R-Anterior-cingulate-G&S_ACC | **-0.04** | 0.05 | 0.98 | **-0.07** | 0.19 | 0.78 | **-0.02** | 0.03 | 0.77 |
| L-Central-S | **-0.04** | 0.04 | 0.98 | **-0.04** | 0.18 | 0.78 | **-0.01** | 0.03 | 0.77 |
| R-Middle-occipital-G | **-0.04** | 0.04 | 0.98 | **-0.04** | 0.18 | 0.78 | **-0.01** | 0.03 | 0.77 |
| R-Transverse-frontopolar-G&S | **-0.04** | 0.05 | 0.98 | **-0.07** | 0.18 | 0.78 | **-0.02** | 0.03 | 0.77 |
| R-Putamen | **-0.05** | 0.05 | 0.98 | **-0.07** | 0.21 | 0.78 | **-0.02** | 0.04 | 0.77 |
| L-Subcentral-G&S | **-0.05** | 0.06 | 0.98 | **-0.08** | 0.22 | 0.78 | **-0.03** | 0.04 | 0.77 |
| Brain-Stem | **-0.05** | 0.06 | 0.98 | **-0.08** | 0.20 | 0.78 | **-0.03** | 0.04 | 0.77 |
| L-parahippocampal-medial-occipitotemporal-G | **-0.05** | 0.05 | 0.98 | **-0.04** | 0.21 | 0.78 | **-0.02** | 0.04 | 0.77 |
| Verbal Comprehension | **-0.05** | 0.06 | 0.98 | **-0.07** | 0.21 | 0.78 | **-0.03** | 0.04 | 0.77 |
| L-Cerebellum-Cortex | **-0.05** | 0.06 | 0.98 | **-0.09** | 0.22 | 0.78 | **-0.03** | 0.05 | 0.77 |
| L-Marginal-cingulate-S | **-0.06** | 0.05 | 0.98 | **-0.05** | 0.23 | 0.78 | **-0.02** | 0.04 | 0.77 |
| R-Accumbens-area | **-0.06** | 0.05 | 0.98 | **-0.07** | 0.21 | 0.78 | **-0.02** | 0.04 | 0.77 |
| R-Planum-temporale-superior-temporal-G | **-0.06** | 0.05 | 0.97 | **-0.03** | 0.21 | 0.78 | **-0.02** | 0.04 | 0.77 |
| L-Superior-frontal-G | **-0.06** | 0.05 | 0.98 | **-0.03** | 0.22 | 0.78 | **-0.02** | 0.04 | 0.77 |
| R-Posterior-lateral-S | **-0.06** | 0.04 | 0.97 | **0.00** | 0.19 | 0.78 | **-0.01** | 0.03 | 0.77 |
| R-Middle-frontal-G | **-0.07** | 0.07 | 0.97 | **-0.06** | 0.26 | 0.78 | **-0.03** | 0.06 | 0.77 |
| L-Hippocampus | **-0.07** | 0.05 | 0.96 | **0.00** | 0.22 | 0.78 | **-0.02** | 0.04 | 0.77 |
| L-Posterior-ventral-cingulate-G vPCC | **-0.07** | 0.04 | 0.96 | **0.05** | 0.18 | 0.78 | **-0.01** | 0.03 | 0.77 |
| R-Postcentral-S | **-0.07** | 0.06 | 0.96 | **-0.01** | 0.23 | 0.78 | **-0.02** | 0.05 | 0.77 |
| R-Superior-occipital-G | **-0.07** | 0.05 | 0.96 | **0.02** | 0.21 | 0.78 | **-0.01** | 0.04 | 0.77 |
| L-Intraparietal-S&transverse-parietal-S | **-0.07** | 0.05 | 0.96 | **0.00** | 0.23 | 0.78 | **-0.02** | 0.05 | 0.77 |
| L-Middle-frontal-G | **-0.07** | 0.07 | 0.96 | **-0.04** | 0.25 | 0.78 | **-0.03** | 0.06 | 0.77 |
| L-Opercular-inferior-frontal-G | **-0.08** | 0.05 | 0.96 | **0.04** | 0.22 | 0.78 | **-0.01** | 0.04 | 0.77 |
| 16PF_Self-Control | **-0.08** | 0.05 | 0.96 | **0.04** | 0.22 | 0.78 | **-0.01** | 0.04 | 0.77 |
| MSCEIT_Experiential | **-0.08** | 0.06 | 0.96 | **-0.01** | 0.25 | 0.78 | **-0.03** | 0.06 | 0.77 |
| R-Superior-frontal-S | **-0.08** | 0.05 | 0.94 | **0.03** | 0.23 | 0.78 | **-0.02** | 0.05 | 0.77 |
| R-Inferior-circular-S-insula | **-0.08** | 0.06 | 0.93 | **0.03** | 0.24 | 0.78 | **-0.02** | 0.05 | 0.77 |
| R-Occipital-pole | **-0.08** | 0.06 | 0.93 | **0.03** | 0.22 | 0.78 | **-0.02** | 0.05 | 0.77 |
| L-Medial-occipitotemporal-S&lingual-S | **-0.08** | 0.05 | 0.93 | **0.04** | 0.20 | 0.78 | **-0.01** | 0.04 | 0.77 |
| L-Inferior-frontal-S | **-0.09** | 0.04 | 0.93 | **0.08** | 0.18 | 0.78 | **-0.01** | 0.04 | 0.77 |
| R-Transverse-temporal-S | **-0.09** | 0.12 | 0.93 | **-0.07** | 0.35 | 0.78 | **-0.07** | 0.15 | 0.89 |
| R-Parieto-occipital-S | **-0.09** | 0.06 | 0.93 | **0.04** | 0.25 | 0.78 | **-0.02** | 0.06 | 0.77 |
| R-Superior-parietal-lobule | **-0.09** | 0.06 | 0.93 | **0.04** | 0.23 | 0.78 | **-0.02** | 0.06 | 0.77 |
| L-Posterior-transverse-collateral-S | **-0.09** | 0.05 | 0.92 | **0.06** | 0.21 | 0.78 | **-0.01** | 0.05 | 0.77 |
| R-Precentral-G | **-0.10** | 0.05 | 0.92 | **0.08** | 0.20 | 0.78 | **-0.01** | 0.05 | 0.77 |
| R-Angular-G | **-0.10** | 0.07 | 0.92 | **0.00** | 0.27 | 0.78 | **-0.03** | 0.07 | 0.77 |
| R-Orbital-G | **-0.10** | 0.05 | 0.92 | **0.07** | 0.22 | 0.78 | **-0.01** | 0.05 | 0.77 |
| L-Superior-occipital-S&transverse-occipital-S | **-0.10** | 0.05 | 0.92 | **0.07** | 0.20 | 0.78 | **-0.01** | 0.04 | 0.77 |
| R-Middle-frontal-S | **-0.10** | 0.05 | 0.92 | **0.06** | 0.22 | 0.78 | **-0.01** | 0.05 | 0.77 |
| L-Transverse-temporal-S | **-0.10** | 0.04 | 0.92 | **0.08** | 0.19 | 0.78 | **-0.01** | 0.04 | 0.77 |
| L-Orbital-G | **-0.10** | 0.05 | 0.92 | **0.08** | 0.21 | 0.78 | **-0.01** | 0.05 | 0.77 |
| R-Cerebellum-Cortex | **-0.10** | 0.06 | 0.92 | **0.02** | 0.25 | 0.78 | **-0.03** | 0.07 | 0.77 |
| L-Lateral-occipitotemporal-S | **-0.11** | 0.05 | 0.91 | **0.09** | 0.21 | 0.78 | **-0.01** | 0.05 | 0.77 |
| R-Middle-anterior-cingulate-G&S_aMCC | **-0.11** | 0.05 | 0.89 | **0.06** | 0.22 | 0.78 | **-0.02** | 0.06 | 0.77 |
| R-Anterior-transverse-collateral-S | **-0.11** | 0.07 | 0.89 | **0.08** | 0.27 | 0.78 | **-0.02** | 0.08 | 0.77 |
| R-Horizontal-anterior-lateral-S | **-0.11** | 0.05 | 0.89 | **0.13** | 0.20 | 0.78 | **0.00** | 0.05 | 0.77 |
| L-Anterior-cingulate-G&S_ACC | **-0.11** | 0.05 | 0.89 | **0.09** | 0.23 | 0.78 | **-0.01** | 0.06 | 0.77 |
| R-Intraparietal-S&transverse-parietal-S | **-0.11** | 0.05 | 0.89 | **0.10** | 0.20 | 0.78 | **-0.01** | 0.05 | 0.77 |
| R-Long-insular-G&central-insula-S | **-0.11** | 0.05 | 0.89 | **0.11** | 0.20 | 0.78 | **-0.01** | 0.05 | 0.77 |
| R-Subcentral-G&S | **-0.11** | 0.05 | 0.89 | **0.11** | 0.20 | 0.78 | **-0.01** | 0.05 | 0.77 |
| R-Marginal-cingulate-S | **-0.11** | 0.07 | 0.89 | **0.09** | 0.25 | 0.78 | **-0.02** | 0.07 | 0.77 |
| L-Superior-circular-S-insula | **-0.11** | 0.05 | 0.89 | **0.08** | 0.23 | 0.78 | **-0.01** | 0.06 | 0.77 |
| R-Temporal-pole | **-0.11** | 0.05 | 0.89 | **0.11** | 0.22 | 0.78 | **-0.01** | 0.06 | 0.77 |
| L-Horizontal-anterior-lateral-S | **-0.12** | 0.05 | 0.89 | **0.07** | 0.20 | 0.78 | **-0.01** | 0.05 | 0.77 |
| L-Short-insular-G | **-0.12** | 0.06 | 0.89 | **0.10** | 0.23 | 0.78 | **-0.01** | 0.06 | 0.77 |
| L-Inferior-precentral-S | **-0.12** | 0.05 | 0.89 | **0.10** | 0.20 | 0.78 | **-0.01** | 0.05 | 0.77 |
| Beck Depression Inventory | **-0.12** | 0.06 | 0.89 | **0.09** | 0.24 | 0.78 | **-0.01** | 0.07 | 0.77 |
| R-Inferior-occipital-G&S | **-0.12** | 0.05 | 0.89 | **0.09** | 0.20 | 0.78 | **-0.01** | 0.06 | 0.77 |
| R-Subcallosal-G | **-0.12** | 0.06 | 0.89 | **0.07** | 0.23 | 0.78 | **-0.02** | 0.06 | 0.77 |
| L-Cuneus | **-0.12** | 0.06 | 0.89 | **0.07** | 0.25 | 0.78 | **-0.02** | 0.07 | 0.77 |
| L-Orbital-S | **-0.12** | 0.05 | 0.89 | **0.12** | 0.19 | 0.78 | **0.00** | 0.05 | 0.77 |
| R-Superior-circular-S-insula | **-0.12** | 0.06 | 0.89 | **0.10** | 0.23 | 0.78 | **-0.01** | 0.06 | 0.77 |
| L-Superior-parietal-lobule | **-0.12** | 0.05 | 0.89 | **0.09** | 0.22 | 0.78 | **-0.01** | 0.07 | 0.77 |
| R-Inferior-precentral-S | **-0.12** | 0.04 | 0.89 | **0.13** | 0.17 | 0.78 | **0.00** | 0.05 | 0.77 |
| L-Postcentral-S | **-0.13** | 0.07 | 0.89 | **0.10** | 0.26 | 0.78 | **-0.02** | 0.08 | 0.77 |
| L-Thalamus-Proper | **-0.13** | 0.05 | 0.89 | **0.11** | 0.20 | 0.78 | **-0.01** | 0.06 | 0.77 |
| L-Orbital-inferior-frontal-G | **-0.13** | 0.06 | 0.89 | **0.10** | 0.25 | 0.78 | **-0.01** | 0.07 | 0.77 |
| L-Transverse-frontopolar-G&S | **-0.13** | 0.05 | 0.89 | **0.13** | 0.21 | 0.78 | **0.00** | 0.06 | 0.77 |
| L-Precentral-G | **-0.14** | 0.05 | 0.89 | **0.14** | 0.21 | 0.78 | **0.00** | 0.07 | 0.77 |
| R-Posterior-transverse-collateral-S | **-0.14** | 0.05 | 0.89 | **0.13** | 0.19 | 0.78 | **0.00** | 0.06 | 0.77 |
| R-parahippocampal-medial-occipitotemporal-G | **-0.14** | 0.05 | 0.89 | **0.14** | 0.19 | 0.78 | **0.00** | 0.06 | 0.77 |
| R-Medial-occipitotemporal-S&lingual-S | **-0.14** | 0.06 | 0.89 | **0.11** | 0.23 | 0.78 | **-0.01** | 0.07 | 0.77 |
| R-Calcarine-S | **-0.14** | 0.05 | 0.89 | **0.13** | 0.20 | 0.78 | **0.00** | 0.06 | 0.77 |
| L-Lateral-occipitotemporal-G | **-0.14** | 0.05 | 0.89 | **0.13** | 0.21 | 0.78 | **0.00** | 0.07 | 0.77 |
| R-Orbital-S | **-0.14** | 0.06 | 0.89 | **0.12** | 0.23 | 0.78 | **-0.01** | 0.08 | 0.77 |
| L-Superior-frontal-S | **-0.15** | 0.05 | 0.89 | **0.14** | 0.19 | 0.78 | **0.00** | 0.06 | 0.77 |
| L-Vertical-anterior-lateral-S | **-0.15** | 0.07 | 0.89 | **0.10** | 0.27 | 0.78 | **-0.02** | 0.10 | 0.77 |
| L-Suborbital-S | **-0.15** | 0.04 | 0.89 | **0.14** | 0.17 | 0.78 | **0.01** | 0.05 | 0.77 |
| R-Paracentral-lobule&S | **-0.15** | 0.05 | 0.89 | **0.15** | 0.18 | 0.78 | **0.01** | 0.06 | 0.77 |
| 16PF_Tough-Mindedness | **-0.15** | 0.04 | 0.89 | **0.16** | 0.19 | 0.78 | **0.01** | 0.06 | 0.77 |
| R-Straight-G | **-0.15** | 0.05 | 0.89 | **0.15** | 0.21 | 0.78 | **0.00** | 0.07 | 0.77 |
| L-Parieto-occipital-S | **-0.15** | 0.05 | 0.89 | **0.16** | 0.20 | 0.78 | **0.01** | 0.07 | 0.77 |
| R-Middle-occipital-S&lunatus-S | **-0.16** | 0.05 | 0.89 | **0.14** | 0.20 | 0.78 | **0.00** | 0.07 | 0.77 |
| L-Ligual-medial-occipitotemporal-G | **-0.16** | 0.05 | 0.89 | **0.15** | 0.21 | 0.78 | **0.00** | 0.07 | 0.77 |
| R-Opercular-inferior-frontal-G | **-0.16** | 0.04 | 0.89 | **0.16** | 0.18 | 0.78 | **0.01** | 0.06 | 0.77 |
| R-Thalamus-Proper | **-0.16** | 0.06 | 0.89 | **0.15** | 0.23 | 0.78 | **0.00** | 0.08 | 0.77 |
| 16PF_Anxiety | **-0.16** | 0.05 | 0.89 | **0.14** | 0.20 | 0.78 | **0.00** | 0.07 | 0.77 |
| R-Superior-frontal-G | **-0.16** | 0.05 | 0.89 | **0.16** | 0.20 | 0.78 | **0.00** | 0.07 | 0.77 |
| L-Straight-G | **-0.17** | 0.05 | 0.89 | **0.18** | 0.19 | 0.78 | **0.01** | 0.07 | 0.77 |
| R-Vertical-anterior-lateral-S | **-0.17** | 0.04 | 0.89 | **0.17** | 0.18 | 0.78 | **0.01** | 0.07 | 0.77 |
| L-Triangular-inferior-frontal-G | **-0.17** | 0.05 | 0.89 | **0.18** | 0.21 | 0.78 | **0.01** | 0.08 | 0.77 |
| R-Posterior-ventral-cingulate-G_vPCC | **-0.18** | 0.04 | 0.89 | **0.17** | 0.18 | 0.78 | **0.01** | 0.07 | 0.77 |
| L-Calcarine-S | **-0.18** | 0.05 | 0.89 | **0.18** | 0.20 | 0.78 | **0.01** | 0.08 | 0.77 |
| L-Inferior-circular-S-insula | **-0.18** | 0.04 | 0.89 | **0.18** | 0.17 | 0.78 | **0.02** | 0.06 | 0.77 |
| R-Cuneus | **-0.19** | 0.05 | 0.89 | **0.19** | 0.19 | 0.78 | **0.02** | 0.08 | 0.77 |
| L-Anterior-transverse-collateral-S | **-0.21** | 0.06 | 0.89 | **0.20** | 0.23 | 0.78 | **0.01** | 0.11 | 0.77 |
| Perceived Stress | **-0.22** | 0.04 | 0.89 | **0.22** | 0.18 | 0.78 | **0.03** | 0.08 | 0.77 |
| L-Temporal-pole | **-0.24** | 0.05 | 0.89 | **0.24** | 0.19 | 0.78 | **0.04** | 0.10 | 0.77 |
| STAI-Trait | **-0.26** | 0.04 | 0.79 | **0.25** | 0.18 | 0.78 | **0.05** | 0.10 | 0.77 |
| L-Paracentral-lobule&S | **-0.29** | 0.05 | 0.44 | **0.29** | 0.19 | 0.78 | **0.07** | 0.12 | 0.77 |
| R-Ligual-medial-occipitotemporal-G | **-0.30** | 0.04 | 0.44 | **0.31** | 0.17 | 0.78 | **0.08** | 0.10 | 0.77 |
| L-Inferior-occipital-G&S | **-0.35** | 0.04 | 0.13 | **0.34** | 0.18 | 0.78 | **0.10** | 0.13 | 0.77 |

## Appendix E. Effect size and sample size estimation for every feature.

The Pearson correlation between SNI and every demographic, psychological, and cortical and subcortical GMV feature was computed to estimate effect size. The sample size for detecting the effect of every feature was estimated assuming that only one effect is hypothesized and tested. Abbreviations: L left, R right, G gyrus/gyri, S Sulcus/Sulci.

| Variables | Correlation with SNI | Sample Size Estimation |
| --- | --- | --- |
| **Demographic Characteristics** |  |  |
| Age | -0.0028916 | 938688 |
| Gender | -0.0337893 | 6869 |
| Education | 0.07748887 | 1301 |
| **Psychological measures** |  |  |
| MSCEIT_Experiential | -0.0776123 | 1297 |
| MSCEIT_Strategic | 0.09220126 | 917 |
| 16PF_Extraversion | 0.35402236 | 57 |
| 16PF_Independence | 0.3137471 | 74 |
| 16PF_Tough.Mindedness | -0.1511065 | 338 |
| 16PF_Self.Control | -0.076433 | 1338 |
| 16PF_Anxiety | -0.1600565 | 300 |
| Perceived Stress | -0.2191902 | 157 |
| Beck Depression Inventory | -0.117672 | 561 |
| PANAS_Positive | 0.19642275 | 198 |
| PANAS_Negative | -0.0035564 | 620543 |
| STAI_Trait | -0.2567068 | 113 |
| Empathy Quotient | 0.21784528 | 160 |
| Verbal Comprehension | -0.0550113 | 2588 |
| Perceptual Reasoning | 0.06415813 | 1901 |
| **Cortical GMV in the Left Hemisphere** |  |  |
| L-Frontomarginal-G&S | 0.00918054 | 93120 |
| L-Inferior-occipital-G&S | -0.3494748 | 58 |
| L-Paracentral-lobule&S | -0.2907009 | 87 |
| L-Subcentral-G&S | -0.054361 | 2650 |
| L-Transverse-frontopolar-G&S | -0.1341769 | 430 |
| L-Anterior-cingulate-G&S_ACC | -0.1114413 | 626 |
| L-Middle-anterior-cingulate-G&S_aMCC | 0.06716702 | 1734 |
| L-Middle-posterior-cingulate-G&S_pMCC | 0.0185314 | 22850 |
| L-Posterior-dorsal-cingulate-G_dPCC | 0.14093978 | 389 |
| L-Posterior-ventral-cingulate-G_vPCC | -0.0671571 | 1734 |
| L-Cuneus | -0.1195903 | 543 |
| L-Opercular-inferior-frontal-G | -0.0770589 | 1316 |
| L-Orbital-inferior-frontal-G | -0.1346853 | 427 |
| L-Triangular-inferior-frontal-G | -0.1700251 | 266 |
| L-Middle-frontal-G | -0.0758849 | 1357 |
| L-Superior-frontal-G | -0.0597285 | 2194 |
| L-Long-insular-G&central-insula-S | -0.0173458 | 26081 |
| L-Short-insular-G | -0.1154706 | 583 |
| L-Middle-occipital-G | -0.0384255 | 5310 |
| L-Superior-occipital-G | -0.036754 | 5804 |
| L-Lateral-occipitotemporal-G | -0.1406978 | 391 |
| L-Ligual-medial-occipitotemporal-G | -0.1581672 | 308 |
| L-parahippocampal-medial-occipitotemporal-G | -0.0509749 | 3015 |
| L-Orbital-G | -0.1029906 | 734 |
| L-Angular-G | -0.0166132 | 28432 |
| L-Supramarginal-G | 0.07104795 | 1549 |
| L-Superior-parietal-lobule | -0.1200344 | 539 |
| L-Postcentral-G | -0.0024642 | 1292603 |
| L-Precentral-G | -0.1376735 | 408 |
| L-Precuneus | -0.0176935 | 25066 |
| L-Straight-G | -0.1659726 | 279 |
| L-Subcallosal-G | 0.01509586 | 34436 |
| L-Anterior-transverse-temporal-G | -0.0394119 | 5047 |
| L-Lateral-superior-temporal-G | 0.01605985 | 30426 |
| L-Planum-polare-superior-temporal-G | -0.0360494 | 6034 |
| L-Planum-temporale-superior-temporal-G | 0.18072367 | 234 |
| L-Inferior-temporal-G | -0.0155332 | 32524 |
| L-Middle-temporal-G | 0.18989198 | 212 |
| L-Horizontal-anterior-lateral-S | -0.1133548 | 605 |
| L-Vertical-anterior-lateral-S | -0.14504 | 367 |
| L-Posterior-lateral-S | -0.0224271 | 15599 |
| L-Occipital-pole | 0.02742809 | 10427 |
| L-Temporal-pole | -0.2367171 | 134 |
| L-Calcarine-S | -0.1771533 | 244 |
| L-Central-S | -0.0401729 | 4858 |
| L-Marginal-cingulate-S | -0.0568431 | 2423 |
| L-Anterior-circular-S-insula | 0.17041338 | 264 |
| L-Inferior-circular-S-insula | -0.1812133 | 233 |
| L-Superior-circular-S-insula | -0.1131711 | 607 |
| L-Anterior-transverse-collateral-S | -0.2075977 | 176 |
| L-Posterior-transverse-collateral-S | -0.0949917 | 864 |
| L-Inferior-frontal-S | -0.0847298 | 1087 |
| L-Middle-frontal-S | 0.00542848 | 266342 |
| L-Superior-frontal-S | -0.1462755 | 361 |
| L-S-intermedius-primus | 0.00539593 | 269566 |
| L-Intraparietal-S&transverse-parietal-S | -0.0729457 | 1469 |
| L-Middle-occipital-S&lunatus-S | -0.0078422 | 127618 |
| L-Superior-occipital-S&transverse-occipital-S | -0.097515 | 820 |
| L-Anterior-occipital-S&preoccipital-notch | -0.0322884 | 7523 |
| L-Lateral-occipitotemporal-S | -0.1044436 | 714 |
| L-Medial-occipitotemporal-S&lingual-S | -0.085147 | 1077 |
| L-Lateral-orbital-S | -0.011093 | 63778 |
| L-Medial-orbital-S | 0.06266573 | 1993 |
| L-Orbital-S | -0.1194376 | 544 |
| L-Parieto-occipital-S | -0.1557196 | 318 |
| L-Pericallosal-S | 0.04500085 | 3870 |
| L-Postcentral-S | -0.1276236 | 476 |
| L-Inferior-precentral-S | -0.1208256 | 532 |
| L-Superior-precentral-S | -0.0239827 | 13640 |
| L-Suborbital-S | -0.1456051 | 364 |
| L-Subparietal-S | 0.0143831 | 37934 |
| L-Inferior-temporal-S | 0.03142184 | 7944 |
| L-Superior-temporal-S | -0.0340876 | 6749 |
| L-Transverse-temporal-S | -0.1001802 | 776 |
| **Cortical GMV in the Right Hemisphere** |  |  |
| R-Frontomarginal-G&S | -0.0189623 | 21823 |
| R-Inferior-occipital-G&S | -0.1175469 | 562 |
| R-Paracentral-lobule&S | -0.1487712 | 349 |
| R-Subcentral-G&S | -0.109243 | 652 |
| R-Transverse-frontopolar-G&S | -0.0426377 | 4311 |
| R-Anterior-cingulate-G&S_ACC | -0.0416376 | 4521 |
| R-Middle-anterior-cingulate-G&S_aMCC | -0.1071467 | 678 |
| R-Middle-posterior-cingulate-G&S_pMCC | -0.0207774 | 18175 |
| R-Posterior-dorsal-cingulate-G_dPCC | 0.03291939 | 7237 |
| R-Posterior-ventral-cingulate-G_vPCC | -0.1752414 | 250 |
| R-Cuneus | -0.184783 | 224 |
| R-Opercular-inferior-frontal-G | -0.1580088 | 308 |
| R-Orbital-inferior-frontal-G | 0.11117239 | 629 |
| R-Triangular-inferior-frontal-G | 0.05084247 | 3030 |
| R-Middle-frontal-G | -0.0664817 | 1770 |
| R-Superior-frontal-G | -0.1610238 | 297 |
| R-Long-insular-G&central-insula-S | -0.1106463 | 635 |
| R-Short-insular-G | -0.0287966 | 9459 |
| R-Middle-occipital-G | -0.0465737 | 3613 |
| R-Superior-occipital-G | -0.0709251 | 1554 |
| R-Lateral-occipitotemporal-G | -0.0149231 | 35238 |
| R-Ligual-medial-occipitotemporal-G | -0.3020131 | 80 |
| R-parahippocampal-medial-occipitotemporal-G | -0.1395864 | 397 |
| R-Orbital-G | -0.0972696 | 824 |
| R-Angular-G | -0.0943059 | 877 |
| R-Supramarginal-G | -0.0194015 | 20846 |
| R-Superior-parietal-lobule | -0.0922448 | 917 |
| R-Postcentral-G | -0.0329844 | 7208 |
| R-Precentral-G | -0.0939214 | 884 |
| R-Precuneus | 0.02509895 | 12453 |
| R-Straight-G | -0.1524617 | 332 |
| R-Subcallosal-G | -0.1208282 | 532 |
| R-Anterior-transverse-temporal-G | 0.04679593 | 3578 |
| R-Lateral-superior-temporal-G | 0.02430314 | 13283 |
| R-Planum-polare-superior-temporal-G | 0.01613924 | 30127 |
| R-Planum-temporale-superior-temporal-G | -0.0613178 | 2082 |
| R-Inferior-temporal-G | 0.04527234 | 3824 |
| R-Middle-temporal-G | 0.00820435 | 116599 |
| R-Horizontal-anterior-lateral-S | -0.1103838 | 638 |
| R-Vertical-anterior-lateral-S | -0.1659003 | 279 |
| R-Posterior-lateral-S | -0.0645492 | 1878 |
| R-Occipital-pole | -0.0867708 | 1037 |
| R-Temporal-pole | -0.1154087 | 583 |
| R-Calcarine-S | -0.1395177 | 397 |
| R-Central-S | 0.00131523 | 4537366 |
| R-Marginal-cingulate-S | -0.1111944 | 629 |
| R-Anterior-circular-S-insula | 0.04419833 | 4012 |
| R-Inferior-circular-S-insula | -0.0820806 | 1159 |
| R-Superior-circular-S-insula | -0.1217987 | 523 |
| R-Anterior-transverse-collateral-S | -0.1085695 | 660 |
| R-Posterior-transverse-collateral-S | -0.1379085 | 407 |
| R-Inferior-frontal-S | 0.00657545 | 181527 |
| R-Middle-frontal-S | -0.0997017 | 784 |
| R-Superior-frontal-S | -0.0829986 | 1133 |
| R-S-intermedius-primus | -0.0242404 | 13352 |
| R-Intraparietal-S&transverse-parietal-S | -0.1109044 | 632 |
| R-Middle-occipital-S&lunatus-S | -0.1580173 | 308 |
| R-Superior-occipital-S&transverse-occipital-S | -0.0236655 | 14009 |
| R-Anterior-occipital-S&preoccipital-notch | -0.0035028 | 639706 |
| R-Lateral-occipitotemporal-S | -0.0317386 | 7786 |
| R-Medial-occipitotemporal-S&lingual-S | -0.1417571 | 385 |
| R-Lateral-orbital-S | 0.04932258 | 3220 |
| R-Medial-orbital-S | -0.0138083 | 41159 |
| R-Orbital-S | -0.1452631 | 366 |
| R-Parieto-occipital-S | -0.0881139 | 1005 |
| R-Pericallosal-S | 0.01345474 | 43351 |
| R-Postcentral-S | -0.0719338 | 1511 |
| R-Inferior-precentral-S | -0.1217513 | 524 |
| R-Superior-precentral-S | 0.12734155 | 478 |
| R-Suborbital-S | 0.08156378 | 1174 |
| R-Subparietal-S | 0.01929028 | 21087 |
| R-Inferior-temporal-S | 0.04123692 | 4610 |
| R-Superior-temporal-S | 4.10E-05 | 4675478286 |
| R-Transverse-temporal-S | -0.0764489 | 1337 |
| **Subcortical GMV** |  |  |
| L-Cerebellum-Cortex | -0.0587125 | 2271 |
| L-Thalamus-Proper | -0.1296967 | 461 |
| L-Caudate | 0.15014569 | 342 |
| L-Putamen | -0.0104299 | 72146 |
| L-Pallidum | 0.04398183 | 4052 |
| Brain-Stem | -0.0536815 | 2718 |
| L-Hippocampus | -0.071467 | 1531 |
| L-Amygdala | 0.12002447 | 539 |
| L-Accumbens-area | 0.22622697 | 147 |
| R-Cerebellum-Cortex | -0.1034844 | 727 |
| R-Thalamus-Proper | -0.1606671 | 298 |
| R-Caudate | 0.17319617 | 256 |
| R-Putamen | -0.0491374 | 3245 |
| R-Pallidum | 0.111796 | 622 |
| R-Hippocampus | 0.03552365 | 6214 |
| R-Amygdala | 0.09236651 | 914 |
| R-Accumbens-area | -0.0557185 | 2522 |

## Data statement

All data are available at https://osf.io/zumwt/?view_only=4f11ca10ed5947c1be1ecdea57cfdff3.

## Acknowledgements

We thank Tim Armstrong for helping with data collection, and Dorit Kliemann and Julien Dubois for helpful discussion. Funding: This work was supported by a Silvio O. Conte Center from the National Institute of Mental Health (2P50MH094258).

## Author contributions

CL carried out preregistration, most data processing except for aspects of the MRI data, and drafted the paper; CL and UK performed all data analysis; JMT carried out MRI data collection, MRI data processing, helped with data analysis, and helped drafting parts of the Methods; MG helped with assembling data, preregistration, and carring out a literature review for the introduction and discussion sections of the paper; LP expanded the Social Network Index to assess modes of communication and types of social support, supervised behavioral data collection, and helped with processing of behavioral data; RA initially conceived of project and helped draft the paper; All authors contributed to intellectual discussions on the project, and all authors participated in revisions to finalize the manuscript.

